# Osmotic pressure gradients in *E. coli* biofilms revealed by in-situ sensors

**DOI:** 10.64898/2026.04.02.716217

**Authors:** Wenbo Zhang, Emanuel Schneck, Luca Bertinetti, Cécile M. Bidan, Peter Fratzl

## Abstract

Osmotic pressure has been known to play essential roles in living systems from single cells to complex tissues. However, direct in-situ measurements of osmotic pressures in biosystems have remained challenging, especially in complicated heterogeneous systems in which osmotic pressure gradients could exist and induce directed forces. Bacterial biofilms –– organized communities of bacteria encased in a self-produced extracellular matrix –– are a major mode of bacterial life. It has, however, remained unexplored how the osmotic pressure is distributed in the biofilm and how this distribution contributes to biofilm growth and activity. Here, liposomal nano-sensors are developed for the in-situ mapping of osmotic pressures at an unprecedented microscale resolution in real time using *Escherichia coli.* biofilm as a model system that develops at the surface of a hydrogel containing the nutrients. The measurements reveal osmotic pressure gradients with a radially increasing trend from the inner regions to the outer regions of the biofilm, which is associated with biofilm formation, morphology, and metabolism. The gradients likely contribute to mechanical properties, internal stresses, and nutrient transport. The sensor readouts also show that there is an osmotic pressure difference between the biofilm and the adjacent medium, which may promote biofilm expansion through matrix swelling and bacteria growth *via* water and nutrient uptake from the surroundings. Our novel approach based on in-situ osmotic pressure mapping in a growing biofilm reveals a sophisticated spatial regulation of physical forces, which may inspire new models and approaches in the field of mechanobiology.

## 1. Introduction

Mechanical forces are recognized as important determinants in numerous cellular and developmental processes across different biological scales^1,2^. Biological systems maintain structural integrity, functionality, and resistance to external stresses through a complex balance of internal forces arising from mechanical stresses generated within cells, extracellular matrices (ECM), and tissues^3–6^. Beyond material structure and properties of cells and ECM as well as ATP-driven forces exerted by molecular motors, osmotic pressure represents another fundamental physical principle determining the mechanical state of biological systems. The ability to respond to osmotic stress is evolutionarily conserved across virtually all organisms, highlighting the importance of osmotic regulation for life processes^7–9^. Osmotic pressure differences drive water flux and thereby regulate cellular volume, shape, and function^10^. These pressure-driven flows are essential for processes ranging from single-cell homeostasis to complex tissue development and organ formation^11,12^. In multicellular systems, osmotic pressure contributes to emergent tissue properties and morphogenetic processes^13,14^. For instance, in developing embryos, osmotic pressure has been demonstrated to work alongside other mechanical forces to guide tissue formation^13^. While internal stresses arising from cytoskeletal dynamics and cell-extracellular matrix interactions are widely studied, the internal forces generated by osmotic pressure gradients have remained less explored. At the molecular level, recent studies show that osmotic pressure can lead to the contraction of biomacromolecular fibers like collagen^12^ – a major structural protein in the extracellular matrix –– and to the hydration response of collagen-mimetic peptides^15^, which may contribute to bone strengthening and developing tissue tension during extracellular matrix development. To better understand the role of osmotic pressure in heterogeneous biological systems, it is essential to map its spatiotemporal distribution.

Despite its fundamental role in biological processes, direct measurement of osmotic pressure within living systems has remained a rarely met need^16,17^. Traditional methods for osmotic pressure determination typically involve extracting samples for *ex situ* analysis using colligative methods, which fundamentally disrupts the spatial organization, temporal dynamics, and the delicate original force balance. This limitation has created a knowledge gap in understanding how osmotic pressures are distributed and regulated within intact biological structures. The ability to perform in-situ measurements of osmotic pressure is particularly valuable for understanding heterogenous biological structures and dynamic biological processes, which can reveal previously undetectable osmotic pressure variations that influence development and homeostasis of biological systems.

Of particular interest in this regard are bacterial biofilms. They represent organized, coordinated, and functional bacteria communities encased in a self-produced ECM comprising polysaccharides, proteins, extracellular DNA, and lipids, which creates a highly hydrated environment that responds dynamically to variations in external physicochemical conditions^18^. As a protected growth mode, biofilms enable microbial survival in hostile environments. The biofilm structures are characterized by spatially distinct regions that exhibit heterogeneous gene expression patterns. The appearance of wrinkles suggests that biofilms can be mechanically stressed locally without any external load^19–21^. The intricate structural organization and metabolic complexity give rise to the analogy of biofilms to tissues of higher organisms. Bacterial biofilms can cause significant pathogenic and environmental challenges, driving the need for developing new analytical approaches in biofilm research^22,23^. During biofilm development, physicochemical cues including osmotic pressure actively mediate structural expansion and spatial organization^24–26^, which involves complex thermodynamic coupling and interactions between substrate and the biofilm. Biofilms are known to be heterogeneous based on chemical gradients which can result in differential gene expression within a monospecies biofilm^24,27–29^, but their mechanical heterogeneity is not well understood. In biofilm research, in-situ osmotic pressure measurement is of special interest given the complex spatial organization and heterogeneous microenvironments characteristic of these bacterial communities. Osmotic pressures in a biofilm likely contribute to nutrient transport, waste removal, antibiotic resistance, and structural integrity^25,30,31^. Furthermore, understanding osmotic pressure regulation in biofilms has significant practical implications, potentially informing improved strategies for biofilm control in medical, industrial, and environmental contexts.

Here, to address the need for in-situ osmotic pressure measurements, we have developed liposomal nanosensors capable of providing spatiotemporal maps of osmotic pressure distributions within living bacterial communities. Our previous work developed micron and submicron liposomes loaded with FRET (Förster resonance energy transfer) donor-acceptor dye pairs, which are responsive to osmotic pressures in solution and have tailored responsiveness^16,17^. Building upon this, we here further developed FRET-based osmotic pressure sensors with improved sensitivity, spatial resolution, and biocompatibility that enable unprecedented insights into osmotic landscapes in biofilms and biological tissues. Their nanoscale dimensions allow them to be incorporated in biofilm matrices without disrupting structural integrity. The sensors architecture consists of highly water-soluble FRET dye pairs within a liposomal membrane. When exposed to osmotic pressure, the sensors rapidly release water, whereby the FRET signal is increased. The FRET pair dyes have relatively long excitation and emission wavelengths (Ex 500-650 nm and Em 575-800 nm for donor, Ex 525-700 nm and Em 625-850 nm for acceptor) which is favorable for circumventing interference of biomolecular autofluorescence occurring mainly around 350-500 nm (absorption) and 350-550 nm (emission). This approach enables both high spatial resolution mapping and dynamic temporal monitoring of osmotic gradients as they evolve during biosystem development. Using these nanosensors, we successfully visualize osmotic pressure gradients within *E. coli* biofilms for the first time (see Figure 1a for a schematic illustration). Our measurements reveal previously undetected spatial variations in osmotic pressure that are associated with structural features and cell metabolism states of the biofilm. These findings provide novel insights into the physical mechanisms underlying biofilm formation, maintenance, and resistance to environmental stresses.

**Figure 1.**
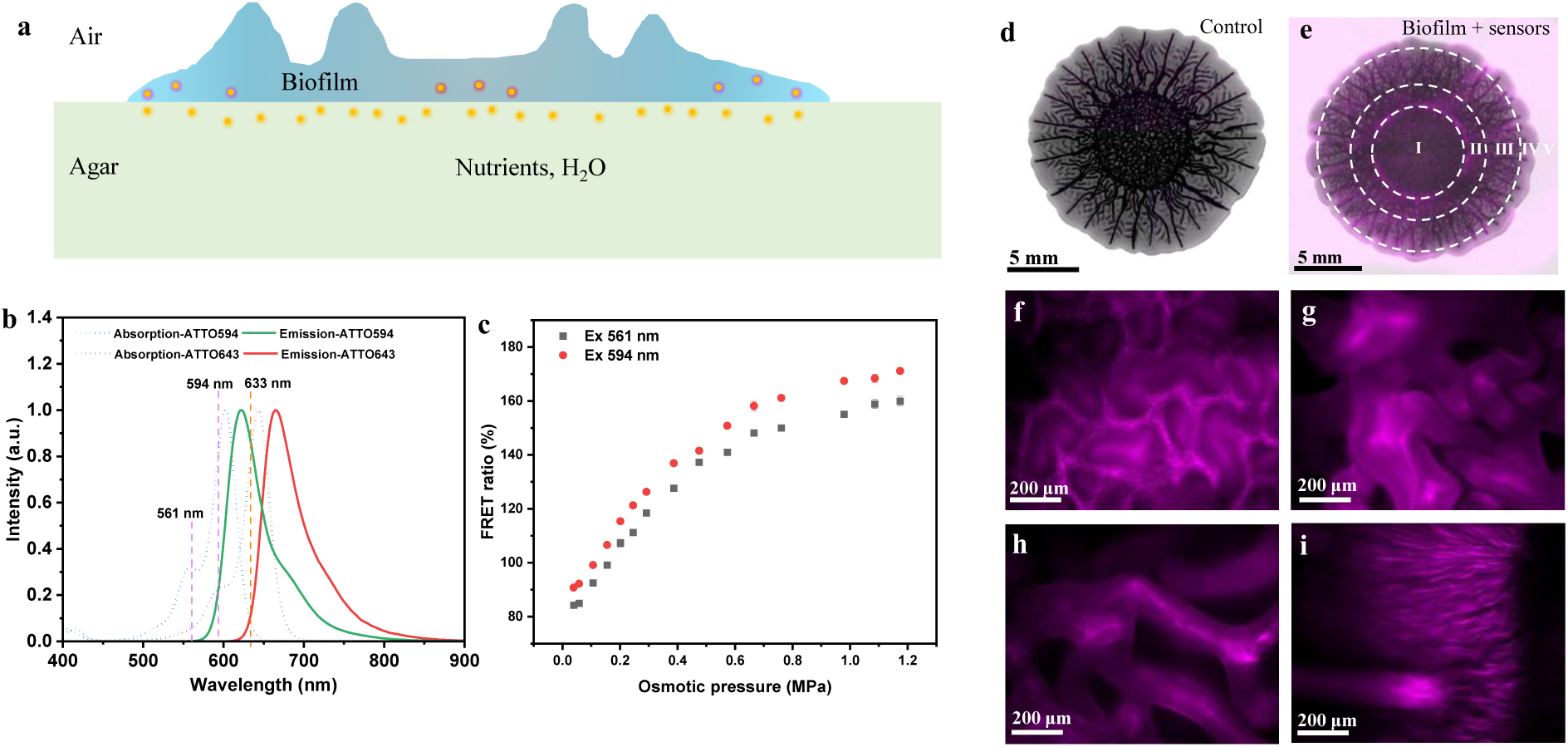
(a) Schematic illustration showing the in-situ sensing of osmotic pressures with liposomal sensors in the biofilm and substrate. The dots in the biofilm and substrate symbolize the liposome sensors with different *R*(П) readouts. Different colors of the dots indicate different readouts of the sensors. (b) Normalized UV/Vis absorption spectra and fluorescence emission spectra of ATTO 594 (donor) and ATTO 643 (acceptor) dyes in water. (c) FRET ratio *R* obtained with the liposomes loaded with a dye concentration of 100 μM (1:1 molar ratio) in 0.05% NaCl as a function of the external osmotic pressure. The excitation wavelength was either 561 nm or 594 nm. (d,e) Wide-field microscopy images of a control *E. coli* biofilm (d) and a biofilm with sensors trapped inside (e) on the substrate. The overlaid fluorescence image in (e) corresponds to the fluorescence of the sensors. (f-i) Fluorescence images at higher magnification of biofilm regions I (f), II (g), III (h) and Ⅳ (i). These regions are indicated in panel (e).

## 2. Results and Discussion

### 2.1. Preparation and Validation of Liposome-based Osmotic Pressure Sensors

The dyes ATTO 594 (MW = 1137 g mol^−1^) and ATTO 643 (MW = 879 g mol^−1^) were chosen as the donor and acceptor fluorophores, respectively, because of their high water solubility and relatively long excitation and emission wavelengths with very limited overlap with biofilm autofluorescence. The fluorescence emission spectra of ATTO 594 and the absorption spectra of ATTO 643 have a large overlap in the in the range of 570–700 nm, which is required for FRET (Figure 1b). The FRET intensity is inversely proportional to the sixth power of the donor/acceptor distance. FRET is therefore extremely sensitive to changes in distance^32,33^, which is the basis for the sensing principle in this study. Great care was taken to determine the ideal excitation wavelength for the FRET pairs, as explained in the following, in order to find the best compromise between minimal donor-acceptor crosstalk, sufficient FRET efficiency, and minimal autofluorescence effect of the biological system.

To study the relation of the FRET efficiency between ATTO 594 and ATTO 643 to their concentration (distance) changes, the fluorescence spectra of their mixed aqueous solutions at a fixed stoichiometry of 1:1 were initially measured for a series of concentrations in the range of 0-100 μM at a fixed excitation wavelength of 514 nm or 561 nm (see section 3.1 in SI for experimental details). As shown in Figure S1a,b, the relative intensity of the acceptor emission peak to the donor emission peak systematically increases with increasing concentration, indicating enhanced sensitized emission of the acceptor at higher concentrations. To quantify the FRET efficiency, the ratio between the emission intensities at 665 nm and 627 nm was defined as the FRET ratio *R*. Figure S1 c shows that *R* increases monotonically with the dye concentration when excited at 514 nm. The solid line in Figure S1c is an empirical fit (coefficient of determination = 0.9999) with *R* = 36.95 + 2.80(μM)^−1^*c* – 0.0113(μM)^−2^*c*^2^, where *c* is the dye concentration. When the excitation wavelength was set to 561 nm, results were similar but with a different function due to the cross-talks and acceptor cross-excitation (Figure S1 d): *R* = 36.69 + 3.04(μM)^−1^*c* – 0.034(μM)^−2^*c*^2^ + 1.39 ×10^−4^(μM)^−3^*c*^3^ (coefficient of determination = 0.9999). However, these effects can be empirically captured by the calibration curves.

For the preparation of the dye-loaded liposomes as the sensors, a dye concentration of either 50 μM or 100 μM (1:1 molar ratio) was used. The osmotic pressure sensors were prepared by an extrusion method (see section 3.2 in SI). The membranes were composed of 1-palmitoyl-2-oleoyl-glycero-3-phosphocholine (POPC) doped with 10% (molar ratio) 1,2-dioleoyl-sn-glycero-3-phosphoethanolamine-N-[methoxy(polyethylene glycol)-2000] (ammonium salt) (DOPE-PEG2000). The liposomes were loaded with a mixed aqueous solution of ATTO 594 (donor) and ATTO 643 (acceptor) dyes in 0.05%wt sodium chloride (NaCl) (8.6 mM). The average hydrodynamic diameter of the liposomes in 0.05% NaCl solution was about 230 nm, as measured by dynamic light scattering (DLS) (Figure S2a). The average zeta potential was ≈-18 mV (Figure S2b). In addition to its use as intra-liposomal osmotic agent, NaCl was also used as extra-liposomal osmotic agent in order to determine the osmotic response of the sensors. To this end, the liposomes were exposed to NaCl solutions of known osmotic strengths and the FRET ratio *R* was determined from the recorded fluorescence spectra. For application of the sensors in biological systems such as biofilms, relatively longer excitation wavelengths are beneficial to limit interferences of biological autofluorescence. However, excitation of the FRET dyes with light of a longer wavelength can result in stronger crosstalk. There is thus a trade-off when choosing the proper excitation wavelength. Therefore, the FRET ratio *R* of the intra-liposomal dyes as a function of the external osmotic pressure Π under excitation with light of different wavelengths was explored (see section 3.4 in SI). Figure S3 shows *R*(Π) in the range 0.04 MPa < Π < 1.2 MPa using a fixed excitation wavelength of 514 nm or 561 nm. Irrespective of the wavelength, *R*(Π) increases monotonically with Π, first approximately linearly at low pressures (≲ 0.6 MPa) and then dependent on pressure more weakly at higher pressures (≳ 0.6 MPa). At the same osmotic pressure, the FRET ratio obtained with 561 nm excitation is slightly higher than that obtained with 514 nm excitation, which can be attributed to stronger crosstalk and cross-excitation. To study whether cross-talk and cross-excitation can affect osmotic pressure sensing, the apparent FRET efficiency *R* was compared to the corrected FRET efficiency *R*_c_ = (*B*_da_ − α*A*_da_ − β*C*_da_)/*A*_da_, where *A*_da_, *B*_da_, and *C*_da_ are the emission intensities at 627 nm, 665 nm, and 700 nm (Excitation 633nm), respectively, of liposomes loaded with the donor/acceptor (50/50 μM), and α = *B*_d_/*A*_d_ and β = *B*_a_/*C*_a_ are correction factors for donor cross-talk and acceptor cross-excitation, respectively. *A*_d_ and *B*_d_ are the emission intensities at 627 nm and 665 nm, respectively, of liposomes loaded with donor dyes only (50 μM); and *B*_a_, and *C*_a_ are the emission intensities at 665 nm and 700 nm (Ex 633 nm), respectively, of liposomes loaded with acceptor dyes only (50 μM). When excited at 514 nm (Figure S4) or 561 nm (Figure S5), *R*_*c*_(Π) exhibits a similar trend as *R*(Π) in the studied range, which verifies that for the sensors with a fixed stoichiometry (1:1) of donor and acceptor, the ratiometric calculation of the apparent FRET efficiency *R* works well for the sensing of osmotic pressures. In the next step, the osmotic response of the sensors was tested in the commonly-used nutrient solutions for bacteria culture at room temperature (r.t.) immediately after addition of the sensors to the solutions. Based on independent measurements with a vapor pressure osmometer (see section 3.3 in SI), the osmotic pressure of a nutrient solution is approximately proportional to the nutrient concentration in the range of interest (Figure S6). As shown in Figures S7 and S8, in solutions containing 2% tryptone/peptone, 1% yeast extract, and additional NaCl (nominal osmolality of NaCl: 0-350 mOsm/kg) (Figure S7) as well as LB broth (1% tryptone, 0.5% yeast extract and 1% NaCl) and dilutions (Figure S8), the osmotic response of the sensors, *R*(Π), is consistent with that in NaCl solutions, demonstrating the functionality of the sensors in the bacteria culture media under the experimental conditions. Moreover, the osmotic response of the sensors remained stable over 72 h at 28 °C. which makes the sensors applicable to monitoring for several days under biofilm formation conditions (Figure S9).

To further explore the feasibility of using a longer excitation wavelength, 594 nm was used to excite the sensors in comparison with 561 nm (Figure 1c), which resulted in similar and consistent relation *R*(Π). Therefore, 594 nm was chosen for the subsequent quantitative imaging studies in the biofilm systems.

### 2.2. Incorporation and Distribution of Osmotic Pressure Sensors in the Biofilm Matrix

To explore the application of the liposomal sensors in biofilms, *Escherichia coli* (*E. coli*) K-12 strain AR3110 was used as a model system. *E. coli* AR3110 forms biofilms by producing both amyloid curli protein and phosphoethanolamine-modified cellulose as major matrix components^31,34,35^. Besides the bacteria and the matrix, biofilms contain more than 80% water^18,31^ and buffer the cells against fluctuations in water availability. Biofilms are characterized by water-filled regions including channels and pores to protect the bacteria against desiccation and support storage and transport of nutrients and waste products^36^. Biofilms can also incorporate material from the surrounding environment, such as minerals and soil particles^37^. Taking advantage of the particle-incorporating ability of the biofilm, the liposomal sensors prepared under sterile conditions were integrated in the biofilms by adding them on top of the agar (1.8%) substrate additionally containing 2% tryptone/peptone, 1% yeast extract but no added NaCl before inoculation of the *E. coli* bacteria (Figure S10) (see sections 3.7.1-3.7.3 in SI). After growing for three days, the biofilms were observed in situ with the wide-field stereomicroscope using the filters for the donor and the acceptor dyes (see section 3.7.4 in SI), respectively (Figure 1d-i; Figure S11,S12). As shown in Figure 1d,e, the biofilm containing sensors (Figure 1d) has a size and morphology similar to that of the control biofilm (Figure 1e) with radial, circumferential, or zigzag wrinkles and delaminated buckles in different regions, which indicates that the incorporation of sensors does not disturb the morphology development during biofilm formation.

The sensors are incorporated relatively evenly throughout the whole biofilm (Figure 1e) as well as in the surrounding agar substrate regions, which is indicated by the fluorescence signal of the sensors in the donor (Figure S11c) and the acceptor channels (Figure 1e, Figure S11d). In contrast, the control biofilm exhibits very limited autofluorescence in both the donor and the acceptor channels (Figure 1d; Figure S11a,b), which minimizes interference of biofilm autofluorescence and sensor fluorescence. Under higher magnification, the sensors are seen to be distributed evenly in different structures of the biofilm in the center region (region I, Figure 1f, Figure S12a), outer region (region II, Figure 1g,h), periphery region (region III, Figure 1i, Figure S12b) and the edge region (region Ⅳ, Figure S12c), as well as in the surrounding substrate (Figure S12d) region. Moreover, due to the passivation of the sensor surfaces with PEG lipids, the sensors are sterically unable to adsorb to the biofilm components^17,38^.

### 2.3. *In situ* 3D Fluorescence Imaging of the Biofilm

In the next step, on the basis of the ex-situ measurement (Figure 1b,c; Figure S1-S9) and the in-situ qualitative studies (Figure 1d-i; Figure S11,S12), the in-situ sensing of osmotic pressures in the *E. coli* AR3110 biofilm-substrate system with the sensors was further explored. Confocal laser scanning microscopy (CLSM) was used for sensitized emission FRET imaging using an excitation wavelength of 594 nm (see sections 3.5 and 3.7.5 in SI). Global preview images (Figure S13) show that the sensor-loaded biofilm and the control biofilm also exhibit comparable morphology, which is consistent with wide-field microscopy observations (Figure 1d,e). The biofilm contains four identifiable regions (Figure S13a, b): Region I, located at the center, is characterized by circumferential corrugated structures. Region II is a transition zone of increasing biofilm thickness with both circumferential and radial wrinkles. Region III features radially orientated ridges that extend toward the periphery of the biofilm. Between these ridges, the biofilm appears essentially flat, as indicated by the homogeneous intensity observed in the transmission image (Figure S13a). Region IV, the outermost zone, is also predominantly flat. The architectural complexity of the *E. coli* biofilm has been proven to be based on three components: the curli fibers, (modified) cellulose, and the bacterial flagellum, where the rings are dependent on the production of both curli and flagella and the axial wrinkles may additionally require the production of cellulose^39–42^. Under the CLSM settings, the observed donor emission (donor channel: Ex 594 nm, Em 604–620 nm) (Figure S13b), sensitized acceptor emission (FRET channel: Ex 594 nm, Em 680–795 nm) (Figure S13c), and direct acceptor emission (Ex 633 nm, Em 680–795 nm) signals (Figure S13d) of the biofilm visualize the presence of the donor, the FRET effect, and the acceptor in the sensors, respectively. The control biofilm has negligible autofluorescence (Figure S13f-h), which is advantageous for the subsequent quantitative FRET imaging.

Then *z*-stacks (150 μm overall *z* range with steps of 10 μm) were collected using CLSM for 3D imaging of the biofilms. Tile scanning was used to image a part of the biofilm from the center to the edge (Figure 2a). Optical sections at a middle plane of the *z* stack show an even distribution of sensors in all biofilm structures (Figure 2b-d). Cross-sectional (*x*, *z*) views of the *z*-stack (Figure 2e) at different positions from the center to the edge (indicated by the green lines on image Figure 2d) reveal that the sensors are incorporated throughout the biofilm from the bottom to the top in all the flat and wrinkled regions. It needs to be noted that higher fluorescence intensity in the FRET channel (sensitized emission) does not necessarily mean higher sensor concentration as the local osmotic pressure can also influence the intensity of sensitized emission. Osmotic pressure sensing is not affected by the sensor concentration because the FRET ratio is independent of the overall intensity.

**Figure 2.**
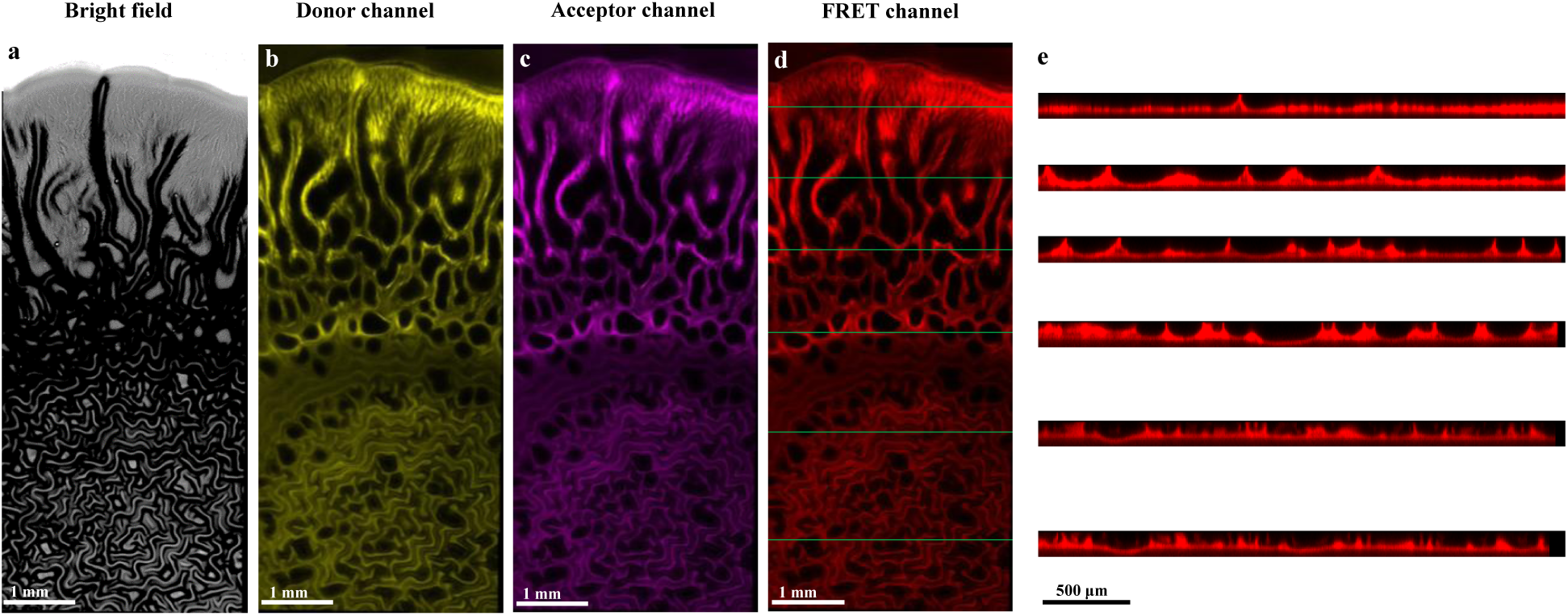
Confocal laser scanning microscopy (CLSM) images of a biofilm with sensors trapped inside. (a) Bright field image of the biofilm. (b) Donor emission signal (Ex 594 nm, Em 604–620 nm). (c) Acceptor emission signal (Ex 633 nm, Em 680–795 nm). (d) Sensitized acceptor emission signal (Ex 594 nm, Em 680–795 nm). (e) X-z cross-sectional view of a z stack (z range 150 μm, step 10 μm) at the positions indicated by the green lines on image (d).

### 2.4. *In situ* Sensing of Osmotic Pressures in the Biofilm

To quantitatively assess the FRET efficiency in the biofilm-substrate system, the FRET ratio *R* is determined for each image sensor pixel emitting sufficiently strong fluorescence signals (see sections 3.5 and 3.7.5 in SI). The fluorescence intensity histograms in the donor channel (Figure S14a,c) and the FRET channel show strong contrast in fluorescence intensity of the sensor-loaded biofilm (Figure S14a,b) and of the control biofilm (Figure S14c,d). By segmentation based on fluorescence intensity, the background pixels and other pixels with insufficient intensity are excluded (threshold intensity: 1200 for the donor channel and 800 for the FRET channel). Figures 3a-e and Video S1 display FRET ratio images of the biofilm at various planes of the *z*-stack from the bottom to the top. The osmotic pressure differences between different regions and planes are clearly identified by their different FRET ratios. The majority of the FRET ratios is in the range of 45%-95% with an average of about 70% (Figure 3f). For quantitative analysis and osmotic pressure mapping, the corresponding *R*(Π) calibration curve was obtained by using the average FRET ratio of the pixels selected by segmentation (excluding pixels with intensities below the threshold value) in solutions containing 2% tryptone/peptone, 1% yeast extract and additional NaCl (nominal osmolality of NaCl: 0-350 mOsm/kg) (see sections 3.6 and 3.7.5 in SI). As shown in Figure S15, *R*(Π) increases systematically with increasing osmotic pressure, from *R* ≈ 47% at Π = 0.05 MPa to *R* ≈ 101% at Π = 0.93 MPa.

**Figure 3.**
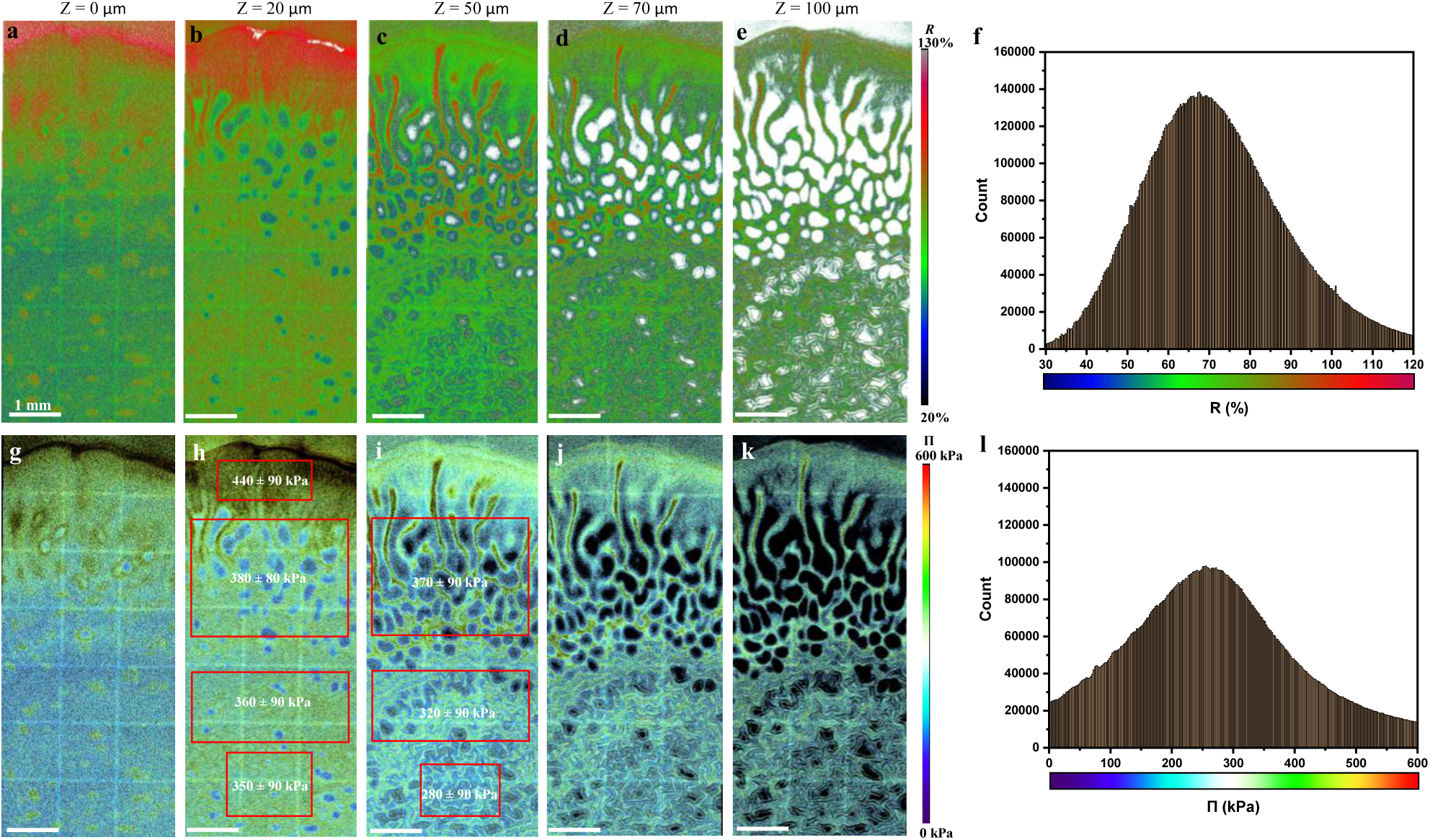
In situ imaging of osmotic pressures in a biofilm. (a-f) FRET imaging of a biofilm region from the center to the periphery with sensors trapped inside at different planes 0 μm (a), 20 μm (b), 50 μm (c), 70 μm (d) and 100 μm (e) of a *z*-stack (z range 150 μm, step 10 μm). (f) Overall FRET ratio distribution in the imaged volume. (g-l) Osmotic pressure mapping of the biofilm at the corresponding z positions: 0 μm (g), 20 μm (h), 50 μm (i), 70 μm (j) and 100 μm (k) of the *z*-stack. The indicated osmotic pressures are average values of the regions in the corresponding red rectangles, which illustrate the osmotic pressure gradients between regions differing in *x*, *y*, and *z*. (l) Overall osmotic pressure distribution in the imaged volume.

The successful *in-situ* osmotic pressure mapping of the biofilm is illustrated in Figure 3g-k and Videos S2. In the osmotic pressure mapping image, the local osmotic pressures can be measured with microscopic spatial resolution. The color-coded measurements revealed a heterogenous osmotic pressure distribution across the different regions. During biofilm formation and expansion, the surface layer of the soft substrate can be deformed^43^ and remodeled^44^ through mechanical forces, nutrient depletion, secretion of metabolites, and biofilm invasion. In fact, it is found that after removal of the AR3110 biofilm from the agar substrate, there is a pattern with depressed and elevated parts in the surface, which conforms to the biofilm morphology. These interactions can lead to physical, chemical, and structural changes in the agar surface, reflecting the dynamic relationship between biofilms and their substrates. During this process, the liposomal sensors can move into the porous substrate layer near the surface^45^ and show signals. As shown in the starting plane image (*z* = 0, Figure 3g), the osmotic pressures in the outer edge region (Region IV) and the outer middle region (Region III) are generally higher than in the inner regions (Regions I, II). Within the inner regions, some localized spots exhibit elevated osmotic pressures compared to their surroundings, which essentially correspond vertically to the “blue hole” areas in the 20 μm plane (Figure 3h). It was reported that biofilms can deform the soft substrate they grow on, where biofilm adhesion transmits buckling-induced stresses to the substrate, generating deformations which turn the initially flat surface into a dome-like shape with elevations in the inner parts and a recess near the biofilm edges^43^. Based on these observations, the areas with higher osmotic pressures in the inner parts in Figure 3g are speculated to be flatter which experienced less stress during biofilm growth which is not enough for buckling. These areas have similar thickness to the flat parts in the periphery regions, as can be differentiated by the relatively light and even grey scales in the transmission image (Figure 2a). The inner regions of the biofilm with concentrated wrinkles are overall elevated, and the substrate surface is also deformed and elevated. In contrast, most of the parts in the periphery regions are flat. In the plane at *z* = 20 μm, the osmotic pressures are higher and more evenly distributed than in the corresponding areas at z = 0. For example, the average osmotic pressure of the rectangle area in the outer edge region is about 440 kPa while in the inner radial wrinkle region it is only about 380 kPa (hole areas are excluded by segmentation). Toward the center, the average osmotic pressure is further decreased: in the outer circumferential wrinkle region it is about 360 kPa and in the center region it is as low as about 350 kPa. In the plane at *z* = 50 μm, most of the flat parts of the biofilm near the edges are close to or beyond the top surface (Figure 3i). In the wrinkle regions where the biofilm is thicker, osmotic pressures in the inner parts are lower than that in the outer parts. For example, the average osmotic pressure of the rectangle area at the center is merely about 280 kPa, in the middle rectangle area is about 320 kPa and in the outer rectangle area is about 370 kPa. Moreover, comparing vertically with the plane at *z* = 20 μm (Figure 3h), the average osmotic pressure in the same horizontal region is found to be decreased. Further up in the planes at *z* = 70 μm (Figure 3j) and *z* = 100 μm (Figure 3k), the wrinkles gradually reach the top from the center to the outer regions where the average osmotic pressures decrease further. Overall, the osmotic pressures are mostly in the range 100 kPa < Π < 400 kPa, with an average of about 270 kPa (Figure 3l).

The 3D-reconstructed *R*/Π images (Figure 4a, b, d, e) and the corresponding cross-section images (Figure 4 c, f) further explicitly demonstrate the 3D distribution of osmotic pressures in the biofilm, echoing the above results and analysis that osmotic pressure gradients exist from the edge to the center and from the bottom to the top. Height dependence plots of the plane-averaged *R* and Π values (Figure 4g, h) show that the average osmotic pressure is highest (≈ 330 kPa) in the range of z = 10-20 μm. In the range of z = 20-50 μm, the plane-averaged osmotic pressure decreases gradually, from ≈ 300 kPa at *z* = 30 μm to ≈ 270 kPa at *z* = 50 μm. In the range of z = 60-100 μm, while the osmotic pressure is still heterogeneously distributed, its plane-averaged value virtually does not change any further and stays around 260 kPa.

**Figure 4.**
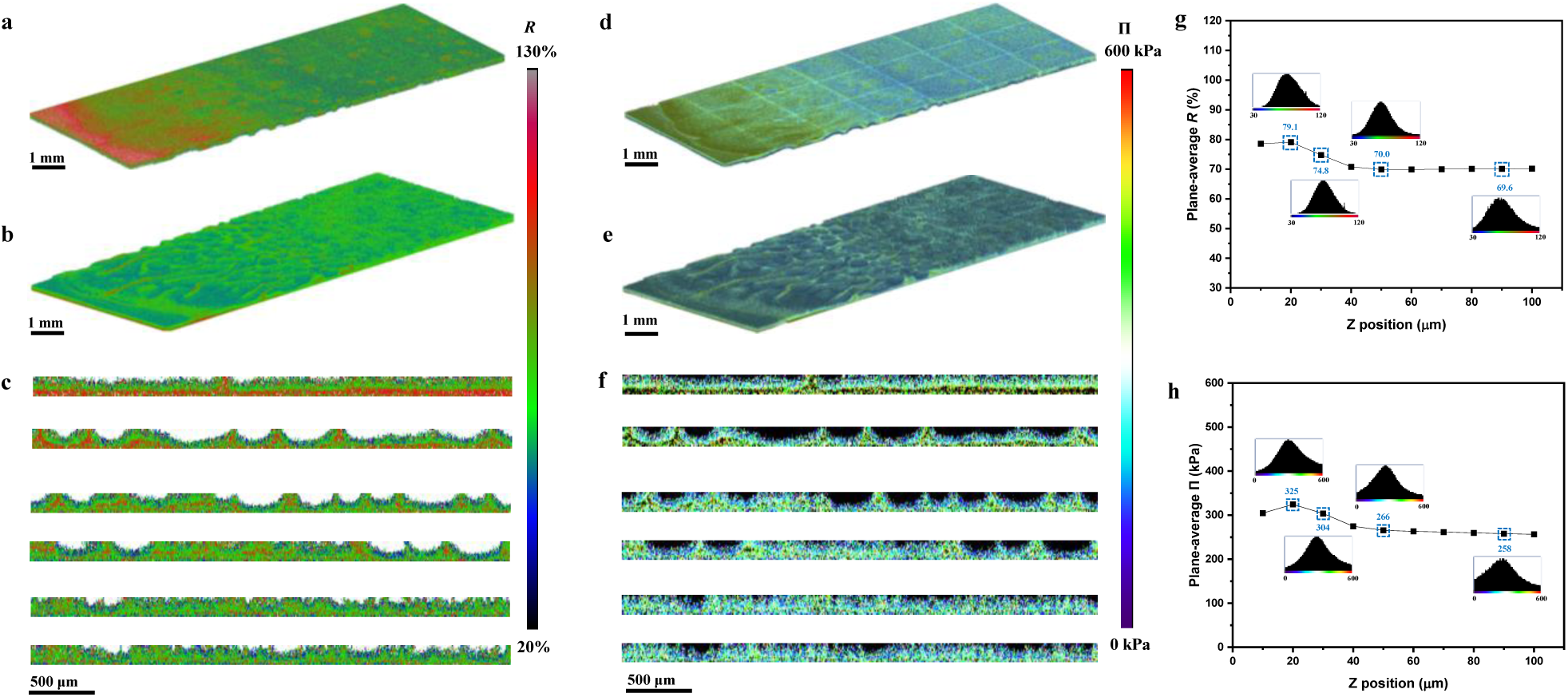
3D distribution of osmotic pressures in the biofilm. (a, b) 3D-reconstructed FRET image (a, facing downward) and a 180-degree rotation (b, facing upward) of a z-stack (z range 120 μm, step 10 μm) of the biofilm with sensors trapped inside. (c) X-z cross-sectional view of the 3D FRET image at the positions indicated by the green lines on image Figure 2(d). (d, e) 3D-reconstructed osmotic pressure image (d, facing downward) and a 180-degree rotation (e, facing upward) of the z-stack. (f) X-z cross-sectional view of the 3D osmotic pressure image at the x-y positions indicated by the green lines on Figure 2 (d). (g, h) Z-profile of the plane-average FRET ratio (g) and osmotic pressure (h) in the z-stack.

To verify the reproducibility of these findings in different biofilms, the osmotic pressure in different regions of matched ROIs and z-stacks from three biofilms grown on the same agar substrate under the same culture conditions were compared. As shown in Figure 5 a-c, the average-intensity projection image of the z-stack of osmotic pressure images of biofilms 1-3 have similar distribution patterns – an overall decreasing trend from the edge region (from around 0 µm, see scale bar on the left of panel a) to the center region (to around 6000 µm) of the biofilms, which is quantitatively demonstrated by the line profile plots of along the longitudinal center line (width 50 pixels) (Figure 5d). Frequency distributions of overall osmotic pressure in the imaged volume of the three biofilms are also comparable (Figure 5e). Quantitative comparison shows that the osmotic pressures of the regions I-IV on images Figure 5a-c and the overall osmotic pressures of the z-stacks of biofilms 1-3 are all closely matched (Figure 5f). The average overall osmotic pressure is 277 ± 5 kPa. Across different regions, the average osmotic pressures exhibit distinct gradients: in region IV (edge region), region II & III (radial wrinkle region and transition region), region I-outer (circumferential wrinkle region-outer) and region I-inner (circumferential wrinkle region-inner) the average values are 331 ± 6 kPa, 280 ± 6 kPa, 261 ± 4 kPa, and 254 ± 3 kPa, respectively. These results demonstrate that the AR3110 biofilms grown under the same conditions have comparable osmotic pressure values and distribution patterns. Moreover, it can be verified that the liposomal sensors and this sensing method work well in different biofilms.

**Figure 5.**
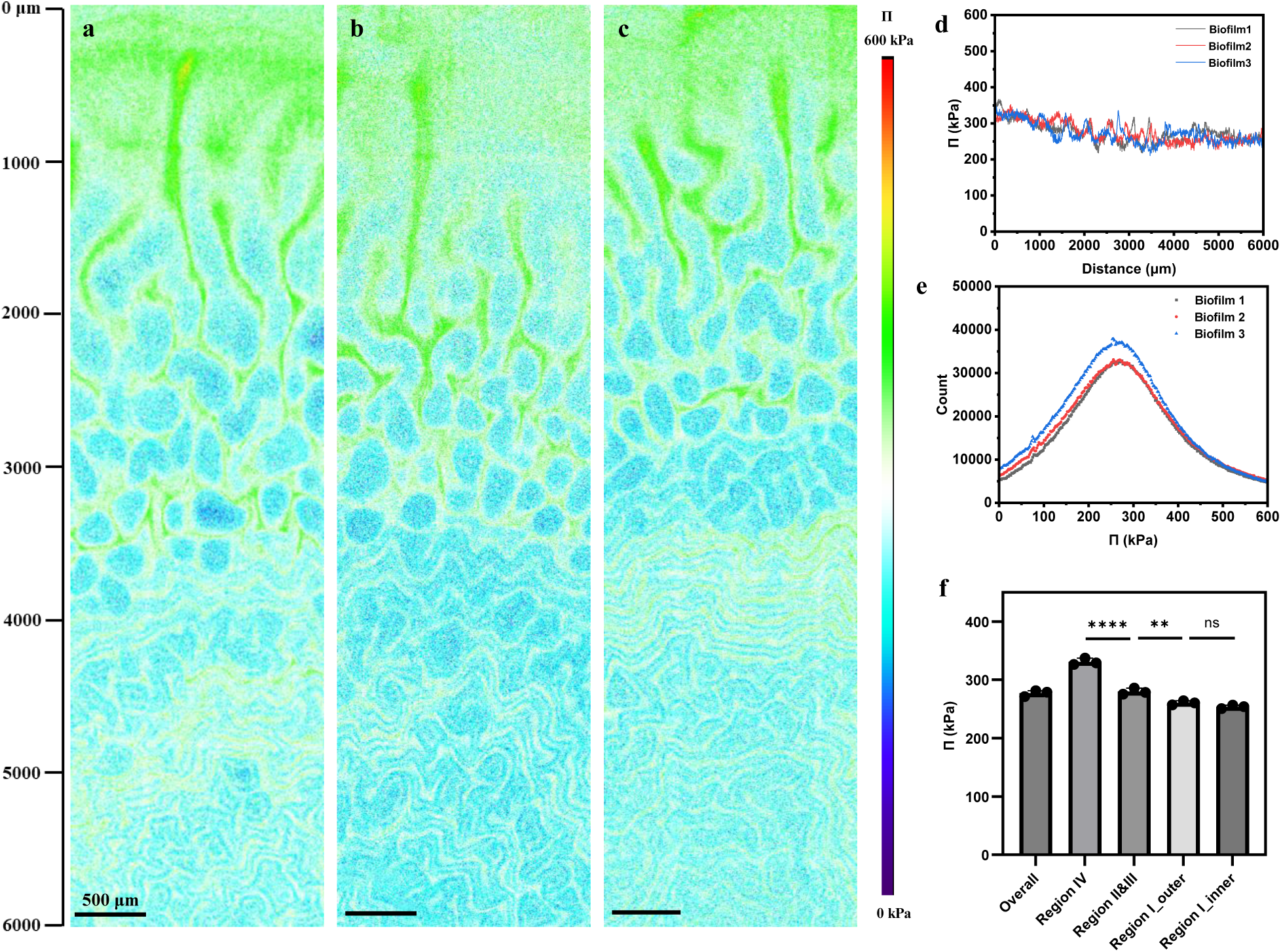
In situ imaging of osmotic pressures in different biofilms. (a-c) Average-intensity projection image of z-stack (z range 100 μm, step 10 μm) osmotic pressure images of biofilms 1-3 with sensors trapped inside. (d) Line profile plots along the longitudinal center line (width 50 pixels) on images (a-c) from the edge region (from around 0 µm, see scale bar on the left of panel a) to the center region (to around 6000 µm) of the biofilms. (e) Frequency distribution of overall osmotic pressure in the imaged volume. (f) Average osmotic pressures of the regions I-IV on images (a-c) and the overall average osmotic pressure of the z-stack of biofilms 1-3. Data are expressed as the mean ± SD, n = 3. ∗∗p < 0.01, ∗∗∗∗p < 0.0001, NS indicates no significant difference at a level of p < 0.05.

The heterogeneous spatial distribution of the osmotic pressures in the biofilm is likely associated with the steady-state evaporation of water. The periphery and the radial ridges surrounded by air are among the more evaporation-dehydrated regions under the influence of factors such as more exposure to air and longer distance from the well-hydrated agar. As a consequence, the osmotic pressure is higher in these regions. In contrast, in the inner regions, the hydrophobic curli molecules form a dense layer on the surface limiting the dehydration^46^. However, the osmotic pressure gradients observed in the biofilm are not solely attributable to the evaporation steady state but also exhibit a significant correlation with the metabolic state of the bacteria within that region (Figure 6). In the biofilm colony, bacterial cells of different states are juxtaposed, and their location is within the different biofilm regions. The bacteria are in a wide range of physiological states with diverse genotypes and phenotypes that express distinct metabolic pathways, stress responses and other specific biological activities^27,47^ (Figure 6a,b). The outer edge of the macrocolony is a growth zone, which is inhabited by long, rod-shaped, and frequently dividing bacteria. These cells are able to produce flagella that get entangled, thereby tying cells together. In the middle and inner zones, the cells in the lower areas nearer to the agar substrate surface are post-exponential, exhibit a shorter rod-shape, and are encased in a dense mesh of flagella filaments. At the upper levels, small ovoid cells surrounded in a dense mesh of curli fibers and cellulose are found, which have entered stationary phase and produce curli and cellulose^40,47^. The formation of this spatial arrangement of physiological differentiation is determined by gradients of nutrients, signaling compounds, and waste products that build up during biofilm growth^27,41^. At the bottom layers and the outer edges of the microcolony, adjacent to the nutrient-rich agar substrate, the cellular morphology indicates active growth physiology. In contrast, bacteria located in the upper regions of the biofilm likely experience nutrient deprivation, primarily due to diffusion limitations and the consumption of available nutrients by actively growing bacteria in the lower layers. Consistently, cells in the upper layers exhibit morphological characteristics indicative of stationary-phase physiology.

**Figure 6.**
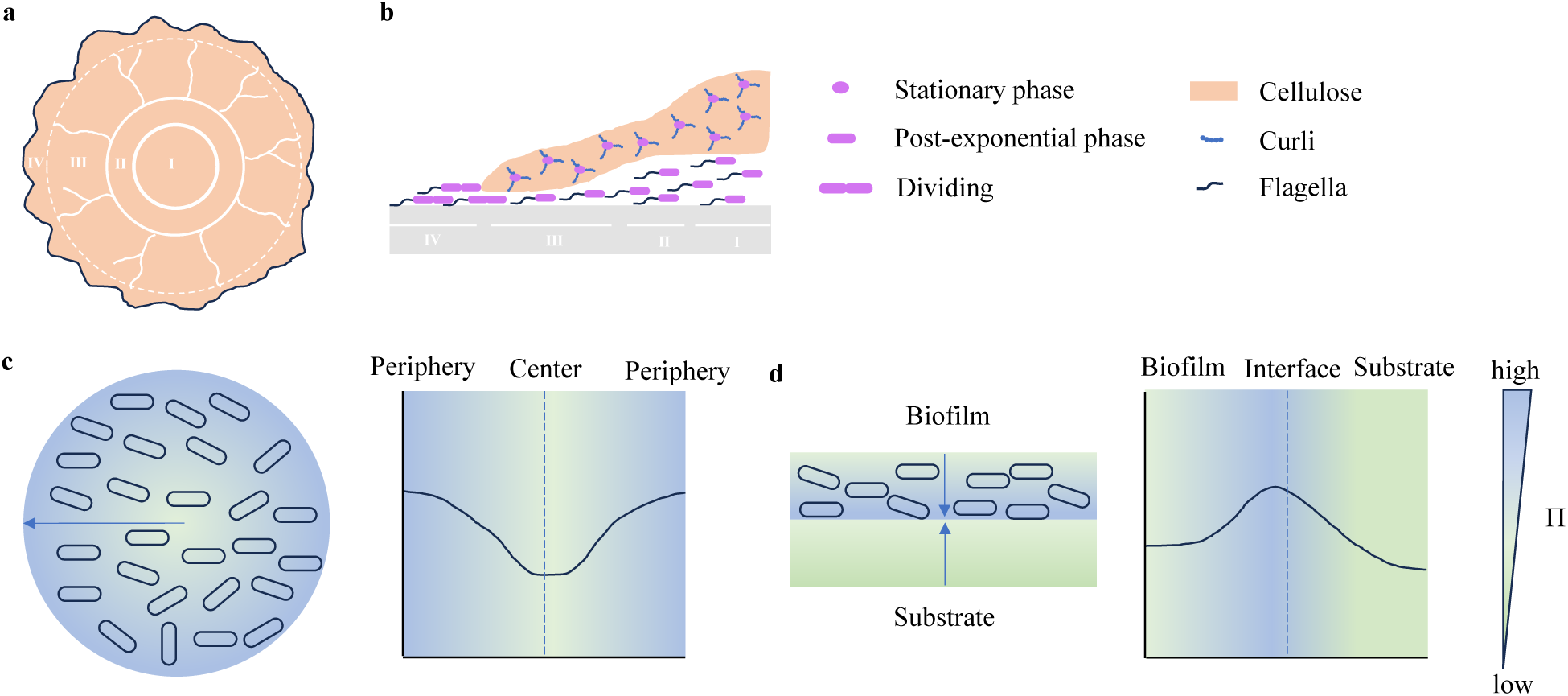
Physiological and matrix composition heterogeneity (a, b), and osmotic pressure gradients in the plane of the biofilm (c) and across its thickness (d). (a) The *E. coli* biofilm structure features concentric rings and axial wrinkles. The rings are dependent on the production of curli and flagella, and the axial wrinkles additionally require the production of cellulose. (b) A cross-section of the colony shows the different cell types co-existing within the biofilm, and their location within the different biofilm regions. (c) Qualitatively distinct pattern of osmotic pressure gradients in a horizontal plane of the biofilm. The osmotic pressures in the outer regions are higher than in the inner regions. (d) Qualitative pattern of osmotic pressure gradients in a cross-section of the biofilm and the substrate near the interface. The osmotic pressures near the interface are higher than those further away from the interface. In (c) and (d), The color gradients indicate osmotic pressure gradients where the blueish and the greenish colors schematically indicate higher and lower osmotic pressure, respectively. The black curves describe the osmotic pressure profiles.

In the locations where the osmotic pressure is higher such as the edge zone of the microcolony and in the lower areas close to the substrate, the bacterial cells are found to be in a metabolically more active state (Figure 6c). When the AR3110 *E. coli* biofilm grows on an agar substrate containing nutrients (2% tryptone/peptone and 1% yeast extract), the distribution of osmotic pressures within the biofilm is collectively influenced by the local components of the substrate and by the biofilm’s metabolic activities. This formation of osmotic pressure gradients appears to be associated with the concentration difference of organic nutrients (such as peptides, amino acids, vitamins, and other growth factors, etc.), salts, metabolic products (such as intermediate products, byproducts, waste products, etc.) and ECM. The overall osmotic environment is a dynamic balance of water retention, nutrient uptake, and the accumulation of solutes within different regions of the biofilm. In the lower layers and periphery of the biofilm, nutrients are more concentrated while the upper layers are more limited in nutrients due to consumption by the bacteria and diffusion limitation. In these regions, the more metabolically active bacteria produce more solute products which can accumulate, particularly when diffusion is limited, leading to localized increases in osmotic pressure.

Besides, it is important to note that this phenomenon is also reflected in the radial wrinkles, which are about several times higher than wide and extend radially toward the periphery of the biofilm. Bacteria within these vertical radial wrinkles keep growing, even some that are located close to the wrinkle tip and are situated at a considerable distance from the nutrient-providing agar. The space at the bottom of the ridges, formed during the upward buckling of the macrocolony at this location, is also entirely filled with actively growing bacteria^40,48,49^. Consistently, the observed osmotic pressures in the interior of the sheath structure of these ridges are higher than in the outer parts of the ridges (Figure 3c-e, i-k), which strongly supports the interpretation that the osmotic pressure is elevated in regions where bacteria are more metabolically active. However, at this point we are unable to provide an answer about what is the cause and what is the consequence of the correlation between high osmotic pressure and high metabolic activity.

It should be noted that osmotic pressure gradients exerted by water evaporation on the biofilm surface can be influenced, in a complex way, by heterogeneity in water mobility within the biofilm. For example, the biofilms have channels of high water mobility, through which the surface would be able to pull water from deeper regions while bypassing the upper layers of the biofilm, which could result in an inversion of the osmotic pressure gradient. Alternatively, the bacteria can consume ATP and pump water inside the biofilm from both the surface and substrate sides, which would also lead to the inverted gradient.

Osmotic pressure gradients can be one of several factors that lead to fluid movement between regions within the biofilm or between the biofilm and its environment. In this dynamic process, water is driven from regions of lower osmotic pressure to regions of higher osmotic pressure to balance the gradient. This steady state water flux affects the hydration state of the biofilm and may also contribute to nutrient transport within it. Moreover, osmotic pressure differences can also affect the mechanical properties of the biofilm and thereby result in expansion, contraction, or buckling. In tendency, osmotic pressure differences drive water influx into regions with higher solute concentrations.

Wrinkles in biofilms play an important role in the colonies’ viability by promoting the transport of nutrients and chemicals, which form as a result of physical, mechanical, and biological processes driven by growth dynamics, nutrient gradients, and mechanical constraints. The differential growth and the spatial constraints imposed by the agar substrate have been identified to lead to mechanical stress within the biofilm which accumulates and eventually causes the biofilm to buckle and form wrinkles as a mechanical response to relieve stress^19,21,47,50^.

The conversion of chemical energy into mechanical energy is a fundamental process in all living systems. While the mechanisms and functions of molecular motors have been well studied^51–54^, the role of osmotic pressure in this context has remained underexplored. Based on our present results, it is presumed that osmotic pressure gradients within the biofilm can also generate internal stresses that promote buckling. These mechanical changes can further modulate nutrient and oxygen distribution, creating feedback loops that influence bacterial metabolism and growth.

Much of the heterogeneity in biological activity within a biofilm can be attributed to microscale variations in solute chemistry^24,27^, with oxygen being the most extensively studied using microelectrodes^55,56^. Our findings indicate that osmotic pressure gradients exist in biofilms grown on agar substrates with nutrients, which has not been reported previously. We hypothesize that these findings can be generalized to other bacterial species as nutrient gradients and physiochemical heterogeneity are often found in the biofilm habitats. This might open up new strategies for biofilm control and contribute to a more holistic view of biofilm formation, development, and evolution.

### 2.5. *In situ* Sensing of Osmotic Pressures in the Substrate near the Interface with Biofilm

The concentration of the solutes and the products in the biofilm may differ from that in the surrounding medium due to selective uptake and retention by the biofilm matrix. The measurement of the osmotic pressure in the substrate is important for understanding the interactions of the biofilm with its environment. The liposomal sensors can move into the mesh-like network with pores of the agar substrate^45,57–59^. Therefore, the osmotic pressures of the substrate layer near the surface can also be measured with the liposomal sensors.

To this end, the substrate under the biofilm was imaged by CLSM using the same settings as that for the biofilm imaging in a z range of 150 μm at a step of 20 μm from the surface to the deeper part (Figure 7). The fluorescence images show the donor emission signal (Figure 7a) and the sensitized acceptor emission signal (Figure 7g), indicating the presence of liposomes in this range of the substrate. The pattern resembling the biofilm morphology in the upper part of the images indicates the deformation of the agar substrate by the biofilm which causes an uneven distribution of the liposomes. The biofilm can physically deform and wrinkle the agar substrate as it grows, creating visible changes in the agar surface due to the forces exerted by the biofilm matrix. The wrinkles are actually vertical deformations of the biofilm together with the adhered substrate^21,43^.

**Figure 7.**
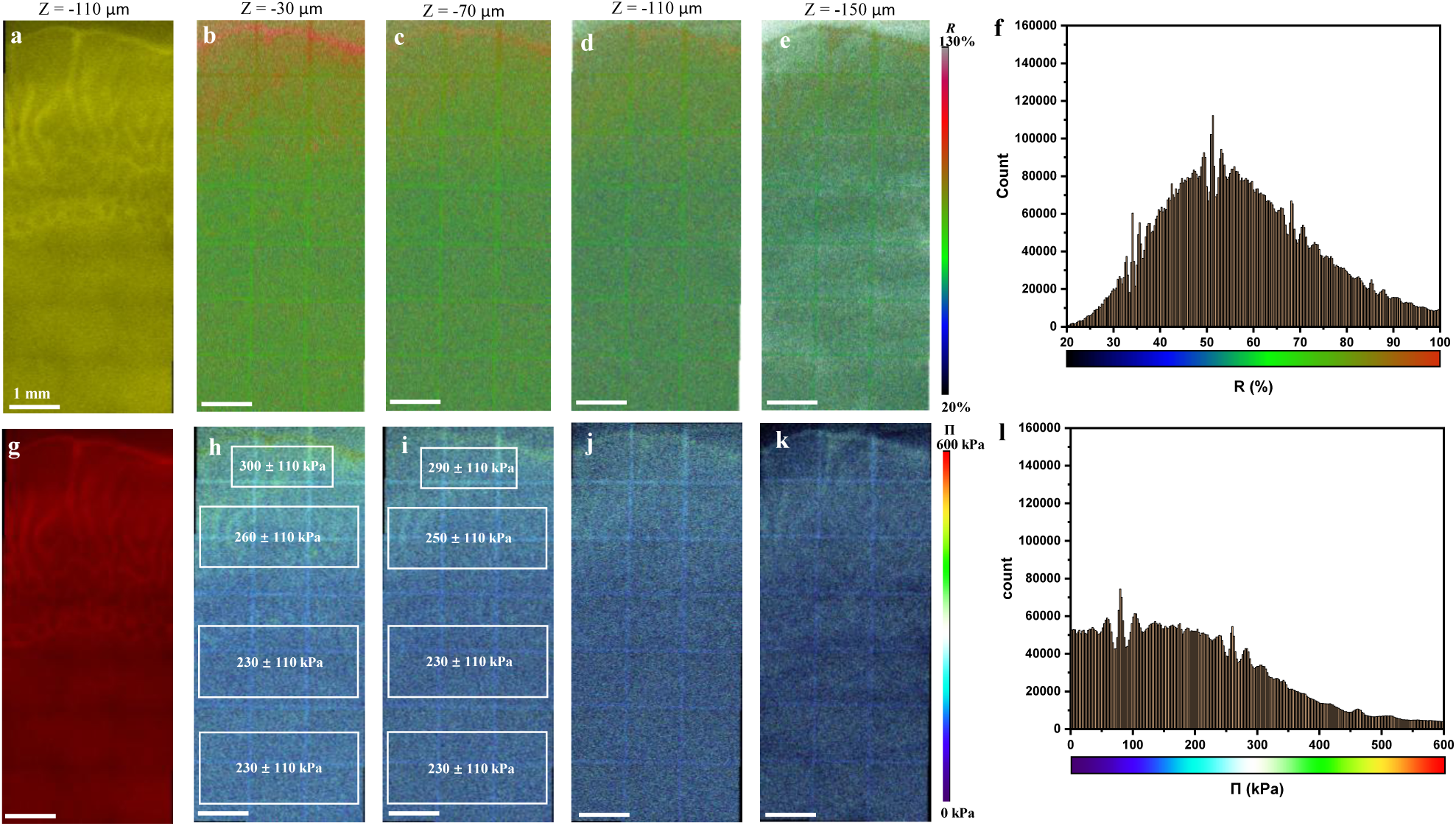
In situ imaging of osmotic pressures in the substrate (z range 150 μm, step 20 μm) under the biofilm. (a) CLSM image showing the donor emission signal (Ex 594 nm, Em 604–620 nm) at plane –110 μm. (b-f) FRET imaging of the substrate at different planes –30 μm (b), –70 μm (c), –110 μm (d) and –150 μm (e). (f) FRET ratio distribution of the z-stack. (g) CLSM image showing the sensitized acceptor emission signal (Ex 594 nm, Em 680–795 nm) at plane –110 μm. (h-l) Osmotic pressure mapping of the substrate at the corresponding z positions: –30 μm (h), –70 μm (i), – 110 μm (j) and –150 μm (k) of the z-stack. (l) Osmotic pressure distribution of the z-stack.

Figures 7b-e and 7h-k exhibit FRET ratio images and corresponding osmotic pressure images of the substrate at various planes of the *z*-stack from the top to the bottom, respectively (see the imaging of the whole substrate z-stack from the bottom to the top in Videos S3,S4). The majority of the FRET ratios of the z-stack are in the range of 35%-80% with an average of about 58% (Figure 7f). Correspondingly, the majority of the osmotic pressures are in the range of 50-300 kPa with an average of about 200 kPa (Figure 7l). Generally, in the same plane in the studied range, the outer regions of the substrate that are under Region III and Region IV of the biofilm have higher average osmotic pressures than the inner regions where the distribution of the osmotic pressure is relatively even. However, the difference between the outer and the inner regions in the same plane decreases with increasing depth below the biofilm. For example, in the plane at *z* = –30 μm (Figure 7h), the average osmotic pressures of the rectangle areas from the periphery to the center are 300, 260, 230 and 230 kPa, respectively. In the plane at *z* = –70 μm (Figure 7i), the average osmotic pressures in the vertically corresponding areas are 290, 250, 230 and 230 kPa, respectively. This is consistent with the osmotic pressure of the original nutrient solution (2% tryptone + 1% yeast extract), which is about 190 kPa as measured by osmometer (Figure S6a), which can be increased as biofilms grow as a result of evaporation and interactions with biofilms. It can to some extent serve as an internal standard to validate the readings of the in situ osmotic pressure sensors.

This distribution of osmotic pressures in the substrate near the biofilm reflects the interplay between the biofilm and the substrate. Because of the proximity of the biofilm, the distribution of osmotic pressures in the top layers of the substrate is affected by the biofilm’s activities. The biofilm can absorb water and consume nutrients from the agar substrate while releasing metabolites. As a result, the in-plane distribution of the osmotic pressures in the top layers of the substrate is correlated with those in the adjacent layers of the biofilm. The further away the substrate layer is from the biofilm, the more homogeneously the osmotic pressure distributed within the layer.

The osmotic pressure in the bottom layer of the biofilm is higher than that in the directly adjacent substrate layer. Based on these experimental results, it is inferred that the biofilm may regulate internal osmotic pressure by accumulating or releasing solutes, which can indirectly affect nutrient uptake from the surrounding environment. Osmotic pressure gradients in the biofilm-substrate system induce fluid flows, increasing transmission of forces through the network and driving enhanced transport of the fluid through the network. The osmotic pressure difference with respect to the substrate is associated with the spreading of the colony^25,26^. It is reported that the growth of the biofilm and the ECM elastic properties can generate internal mechanical stress as a result of geometrical constrains^20^. Our results suggest that osmotic pressure gradients in the biofilm can be an origin of internal stress. The osmotic pressure gradient is likely a mechanism of force regulation within a biofilm that shapes the biofilm, regulates internal stress, and promotes colony expansion and mechanical resistance. Biosystems are engineered by organisms like strengthened edifices combining pre-strain and internal stress. The current reported understanding of the microscopic osmotic pressure distribution in biological systems is mainly based on deduction and model analysis. The direct measurement of osmotic pressure distribution within a heterogeneous biosystem can give new insights into the origin of internal stress and its role in the development, functions as well as interactions with the environment of the biosystem, which may inspire new models and approaches in the field of mechanobiology.

## 3. Conclusions

In situ monitoring of physicochemical parameters in biosystems and understanding their complex interplay require suitable measurement approaches capable of providing spatiotemporal information without disrupting the biosystem’s native behavior. Our novel liposomal nano-sensors address this need by enabling direct visualization of osmotic pressure distributions within living E. coli biofilms, revealing previously undetectable osmotic gradients in correlation with biofilm formation, morphology and metabolism. We found that there is a radially increasing trend of osmotic pressure from the inner regions to the more metabolically active and more evaporation-dehydrated outer regions. The observation of osmotic pressure gradients in the biofilm indicates that these contribute to mechanical properties, internal stresses, and nutrient transport. Moreover, the osmotic pressures in the biofilm are found to be overall higher than in the adjacent substrate, which is correlated with the spreading of the colony. Our approach offers insights into the physical forces shaping biofilm development, behavior, and interactions with the environment, which holds promise for both fundamental research and applied biofilm management strategies. By resolving spatial distributions of osmotic pressures in intact living systems, our findings underscore the sophisticated spatial regulation of physical forces in developing biological systems, which may inspire new models and approaches in the field of mechanobiology.

## Supporting Information

Supporting Information is available from the authors.

## Acknowledgements

We thank Dr. Rumiana Dimova, Carmen Remde, Dr. Ricardo Ziege, Christine Pilz-Allen and Dr. Laura Zorzetto for their support in confocal microscopy, wide-field fluorescence microscopy and bacteria culture. We thank Prof. Dr. Regine Hengge (Humboldt-Universität zu Berlin) for kindly providing the E. coli bacteria. This work was partially supported by the German Research Foundation (DFG) within SFB1444 as well as by the MaxWater initiative of the Max Planck Society.

## Conflict of Interest

The authors declare no conflict of interest.

## Data Availability Statement

The data that support the findings of this study are available from the corresponding authors upon reasonable request.

## Notes

### Competing Interest Statement

The authors have declared no competing interest.

