## Supporting information for "Osmotic pressure gradients in *E. coli* biofilms revealed by in-situ sensors"

### 1. Materials

1-palmitoyl-2-oleoyl-glycero-3-phosphocholine (POPC) and 1,2-dioleoyl-sn-glycero-3-phosphoethanolamine-N-[methoxy(polyethylene glycol)-2000] (ammonium salt) (DOPE-PEG2000) were purchased from Avanti Polar Lipids (Alabaster, USA). ATTO 594 carboxy and ATTO 643 carboxy were purchased from ATTO-TEC GmbH (Siegen, Germany). Sodium chloride (NaCl) was purchased from neoLab Migge (Heidelberg, Germany). Chloroform was purchased from Merck KGaA (Darmstadt, Germany). All chemicals used in the experiments were of pharmaceutical standard and analytical grade. Luria−Bertani (LB) medium, LB agar, agar-agar, tryptone/peptone from casein and yeast extract were purchased from Carl Roth GmbH (Karlsruhe, Germany), which were all for microbiology. The water used in all experiments was ultrapure water (18.2 MΩ).

### 2. Characterization

**Size/Zeta potential** was measured with a size/ zeta potential analyzer (Zetasizer Nano-ZS, Malvern) equipped with a 632.8 nm He-Ne laser at room temperature (25 $^{o}$C). Each value was averaged from three parallel measurements. **Ultraviolet-visible (UV–Vis) absorption spectra** were recorded with an Analytik Jena UV-Vis Specord 210 Plus spectrophotometer. **Fluorescence emission spectra** were recorded with a BioTek Cytation 5 microplate reader (bulk dye solutions) or a Horiba Fluoromax4 spectrofluorometer. For UV-Vis spectra, fluorescence spectra and zeta potential/size measurements, the sample solutions were used directly. **Confocal laser scanning microscopy (CLSM)**: The liposome suspension in NaCl solutions or media on a 20 mm cell culture dish with a glass bottom was observed with a Leica TCS SP8 system (10×/NA 0.3 objective using commercial software).

### 3. Experimental methods

#### 3.1 Emission spectra measurement of bulk dye solutions

The emission spectra of ATTO 594 carboxy and ATTO 643 carboxy 1:1 mixture solutions (3.13-100 μM) were measured with a BioTek Cytation microplate reader. The solutions were measured in a black 96-well plate with a clear glass bottom and a clear lid (Corning CLS4580). The sample volume was 100 μL per well. The excitation wavelength was set at 514 nm or 561 nm. The emission spectra were recorded at 524-700 nm (Ex 514 nm) or 571-700 nm (Ex 561 nm). Bandwidths for the excitation and emission path were both 10 nm. Gain value was 65 (Ex 514 nm) or 60 (Ex 561 nm). The microplate reader uses a front-face geometry for the fluorescence measurement, which is appropriate for relatively high concentration of dyes as used in this experiment.

#### 3.2 Preparation of liposomal sensors

The liposomes were prepared by the extrusion method using a Mini-Extruder (Avanti Polar Lipids Inc.). Removal of extra-liposomal free dyes was achieved via rinsing and centrifugation or gel permeation chromatography. 5 mg POPC and 1.9 mg DOPE-PEG2000 lipids were dissolved in 0.5 mL chloroform and were evaporated by passing a gentle stream of nitrogen over the sample, followed by drying under vacuum for overnight. The lipid film was hydrated with 0.5 mL mixture solution of ATTO 594 carboxy (100 μM) and ATTO 643 carboxy (100 μM) in 0.05% NaCl solution for 1 h at 30 $^{o}$C. Then the mixture was vortexed and was subjected to 5 freeze/thaw cycles by alternately placing the sample vial in a liquid nitrogen bath and warm water bath. The suspension was extruded through a polycarbonate membrane (pore size 1.0 μm; Avanti Polar Lipids, 610010) 21 times at 30 $^{o}$C. The liposomes were incubated at 4 $^{o}$C overnight. Then free dyes were removed by gel permeation chromatography on Sephadex G-50.

#### 3.3 Osmotic strength measurement of standard solutions

The NaCl standard solutions used for testing the osmotic responses of the liposomal sensors to obtain the calibration curves were prepared with the weighing method at room temperature. The osmolalities of NaCl solutions were determined from freezing point depression using an OSMOMAT 3000 Osmometer (Gonotec GmbH). Commercial standards for the osmometer (0, 300, and 850 mOsm/kg) were analyzed prior to samples which were measured at least in duplicate. Osmolalities of LB media or nutrient (tryptone, yeast extract) solutions with additional NaCl (nominal osmolality of NaCl: 0, 50, 100, 150, 200, 300 or 350 mOsm/kg) solutions were determined from vapor pressure depression using a VAPRO MODEL 5600 Osmometer (ELITech Group, Inc.). Commercial standards for the osmometer (100, 290 and 1000 mOsm/kg) were analyzed prior to samples which were measured at least five times.

#### 3.4 Calibration curves measurement of sensors in standard solutions with spectrofluorometer

The liposomal sensors were dispersed in a 0.05% NaCl solution at a concentration of 2.5 mg/mL. Then 5 µL suspension was added to 100 µL of NaCl/LB/nutrient standard solutions and the emission spectra were recorded with spectrofluorometer (Horiba MC Fluoromax4). The excitation wavelength was set at 514 nm, 561 nm or 594 nm and the emission spectra were recorded at 614-800 nm. Bandwidths for the excitation and emission path were both 6 nm. Integration time was 0.1 s. After adding sensor/0.05% NaCl, the osmotic pressure of the standard solutions is lowered because of the dilution with the added 0.05% NaCl. So the osmotic pressure of the mixture was corrected by calculation according to the osmotic pressure-mass fraction calibration curve obtained using the osmometer. The FRET ratio $R$ ($F_{665}/F_{627}$) was calculated and its variation with varying osmotic pressure was analyzed to study the osmotic response of liposomes in different solutions.

For measurements in LB media or nutrient (tryptone, yeast extract) solutions with additional NaCl (nominal osmolality of NaCl: 0, 50, 100, 150, 200, 300 or 350 mOsm/kg), background signals were measured under the same conditions: 5 μL 0.05% NaCl solution was added to 100 μL the standard solutions and the emission spectra were recorded as above. The FRET ratios were calculated by using corresponding fluorescence signals of the liposome sensors obtained by subtracting the background from the total signals.

**3.5 Confocal microscopy image acquisition and analysis**

The sensor suspensions were imaged on a Leica TCS SP8 confocal laser scanning microscope for FRET evaluation using a 10×/NA 0.3 objective. Confocal laser scanning microscopy (CLSM) allows for image acquisition of different channels pixel-per-pixel practically at the same time.

The three channel settings were as follows: donor channel, excitation at 594 nm, detection at 604-620 nm; FRET channel, excitation at 594 nm, detection at 680-795 nm; acceptor channel, excitation at 633 nm, detection at 680-795 nm. All settings (HyD detector parameters (voltage, offset), pixel dwell time, laser power, electronic zoom and pinhole) were kept constant across all FRET experiments. The image sets of the samples were acquired for FRET evaluation. In the liposomes, the donor and acceptor had a fixed 1:1 stoichiometry. Therefore, the ratio of FRET signal (sensitized acceptor emission) intensity $F_{FRET}$ to the donor signal intensity $F_{donor}$ was adopted as the index of FRET efficiency (*R*). The pixel-by-pixel FRET ratio images were obtained by processing the raw image data sets with the Fiji software.[1]

**3.6 Calibration curves measurement of sensors in nutrients with confocal microscopy**

The liposomal sensors loaded with a dye concentration of 100 μM (1:1 molar ratio) were dispersed in 0.05% NaCl at a concentration of 10 mg/mL. Then 2.6 μL suspension was added to 50 μL 0.05% NaCl or nutrient (tryptone, yeast extract) solutions with additional NaCl (nominal osmolality of NaCl: 0, 50, 100, 150, 200, 300 or 350 mOsm/kg). The osmotic pressure of the mixture was corrected by calculating according to the osmotic pressure-mass fraction calibration curve. A drop of each suspension was placed on 20 mm cell culture dish with a glass bottom. The image sets for the donor, FRET and acceptor channels were acquired with confocal microscopy (Leica TCS SP8, 10×/NA 0.3 water objective), as stated above. For each sample, at least three image sets were recorded in different areas of the sensor suspension drop. NaCl or nutrient solutions without sensors as control samples were measured under the same conditions. Using Fiji software, the average FRET ratio was obtained as follows. The fluorescence intensity sum of an ROI (800 μm × 800 μm in the center of the image) in a donor channel or FRET channel image was calculated. The FRET ratios were calculated by using corresponding fluorescence intensities of the liposome sensors obtained by subtracting the background from the total intensities: FRET ratio ($R$) = $F_{FRET}$/ $F_{donor}$. Then the average FRET ratio of the parallel image sets was calculated and was plotted versus osmotic pressure to calibrate the osmotic response of sensors in nutrient (tryptone, yeast extract) solutions with additional NaCl.

#### 3.7 Biofilm experiments

##### 3.7.1 Preparation of agar plates

Salt-free agar plates (15 cm diameter) were prepared with 1.8% w/v of bacteriological grade agar−agar (Roth, 2266), supplemented with 1% w/v tryptone (Roth, 8952) and 0.5% w/v yeast extract (Roth, 2363).

##### 3.7.2 Bacterial strain and growth.

Bacterial strain *E. coli* K-12 AR3110 was used in this study as the biofilm-forming bacterial strain.

To prepare for the biofilm growth, first, the bacteria suspension was prepared from a single bacterial colony and was grown for about 10 h in LB medium at 37 °C with shaking at 250 rpm. Then each agar plate was inoculated with drops of 5 μL bacteria suspension (OD600 ~4.5) in a 3×3 array. After inoculation, the bacteria suspension drops were dried naturally for 1 h at r.t. The biofilms were grown for 5 d at 28 °C under static conditions in the incubator.

#### 3.7.3 Trapping of liposomal sensors in biofilm-agar system

The biofilms with trapped liposomal sensors were grown on an agar plate without additional osmolytes. For the trapping of sensors in the biofilm-agar system, before inoculation, add 25 μL liposomal sensors (10 mg/mL in 0.05% NaCl) and spread evenly in a 2 cm × 2 cm area. For the control group, 25 μL 0.05% NaCl were added instead of the sensor suspension. These areas were in a 3×3 array consisting of five sensors areas, three 0.05% NaCl areas and one blank area. The sensors suspension or 0.05% NaCl on the agar plate were dried naturally at r.t. for 1.5 h. Then a drop of 5 μL bacteria suspension (OD600 ~4.5) was added to the center of each area and was dried naturally for 1 h at r.t. The biofilms were grown for 3 d at 28 °C under static conditions in the incubator.

#### 3.7.4 *In situ* imaging of biofilms by wide-field microscopy

Biofilms on the agar plate in a 15 cm Petri dish were imaged *in situ* with an AxioZoomV.16 stereomicroscope (Zeiss, Germany).

Biofilm-agar system with trapped sensors was observed by using the following filters: donor channel, 20 Rhodamine filter set (Ex BP 546/12 nm, Beamsplitter FT 560nm, Em BP 575-640 nm); acceptor channel, 50 Cy5 filter set (Ex BP 640/30 nm, Beamsplitter FT 660nm, Em BP 690/50 nm).

#### 3.7.5 ***In situ* sensing of osmotic pressures in biofilm-agar system by FRET imaging with the sensors**

The *in-situ* sensing of osmotic pressures in the biofilm-agar system with the sensors was conducted by using the confocal microscopy (Leica TCS SP8, 10×/NA 0.3 objective).

Some pieces of agar substrate from the same plate with biofilms to be imaged were put in a 20 mm culture dish with a glass bottom. The dish was sealed with parafilm and the humidity in the dish chamber equilibrated overnight. A biofilm together with its underlying substrate was transferred to the glass bottom of the culture dish with pieces of agar substrate around. The substrate bottom was closely attached to the glass without bubbles in between. After sealing the dish with parafilm, 3×7 tile regions were imaged over a z range of 150 μm (step of 10 μm) for the biofilm and a z range of 200 μm for the underlying substrate (step of 20 μm). Three biofilms grown on the same agar substrate were imaged.

The image sets for the donor, FRET and acceptor channels were acquired, as stated above. The transmission channel was illuminated by the 633 nm laser line. For the mapping of FRET ratio of the samples, the following steps were carried out. Using Fiji software, by segmentation the sensor pixels are selected and used for the calculation of pixel-by-pixel FRET ratio. After excluding outliers, the FRET mapping was achieved. By taking advantage of the $R$-$\Pi$ calibration curve, osmotic pressures can be quantified spatially on the FRET image.

#### 3.7.6 Statistical analysis

Data are expressed as the mean ± standard deviation (SD). Statistical analysis was determined by one-way analysis of variance (ANOVA) with the Origin software. Means comparison was performed with the Tukey's method. A p value $< 0.05$ is considered statistically significant.

### 4. Supplementary results

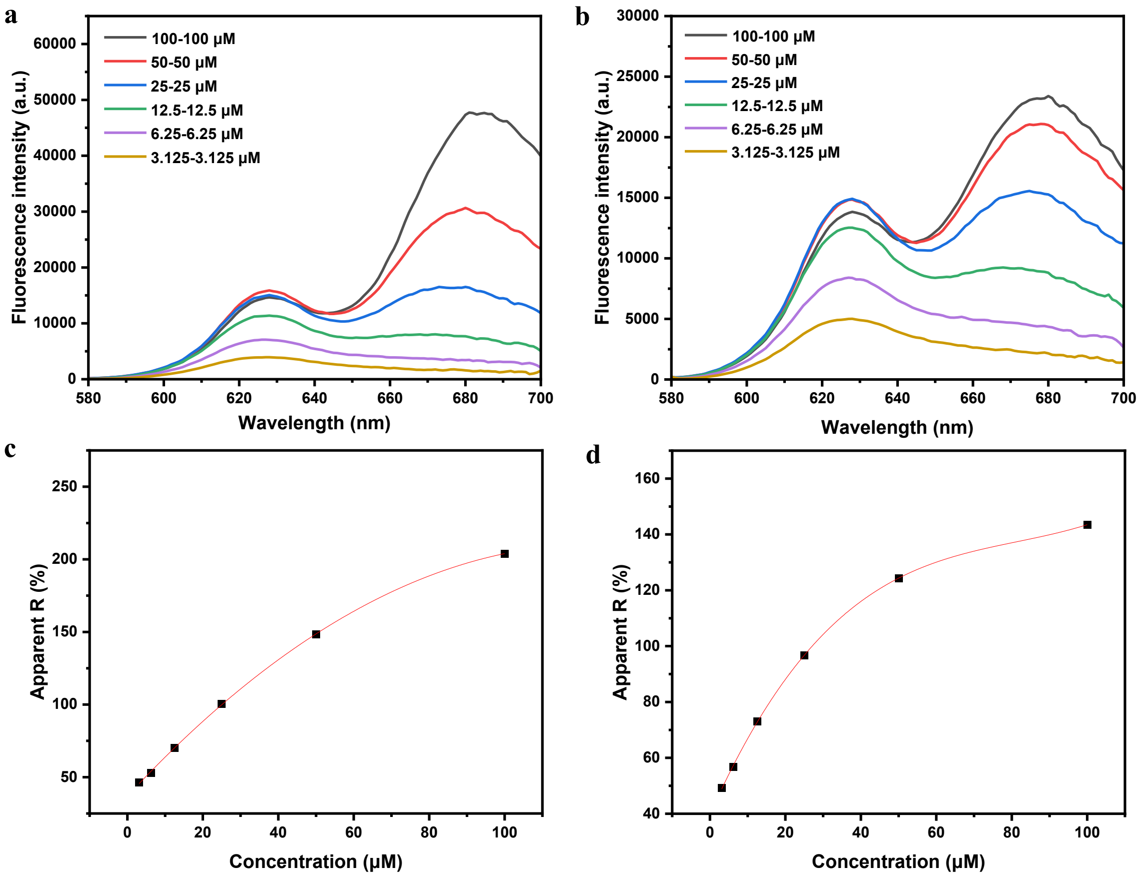


**Figure S1**. (a, b) Fluorescence emission spectra of ATTO 594-ATTO 643 mixture (1:1 molar ratio) in water with different concentrations (3.13–100 μm) at a fixed excitation wavelength of 514 nm or 561 nm. (c, d) Emission ratio *R* as a function of the dye concentration in a 1:1 molar ratio in bulk aqueous solution. The solid line is an empirical fit to the data points (coefficient of determination =0.9999).

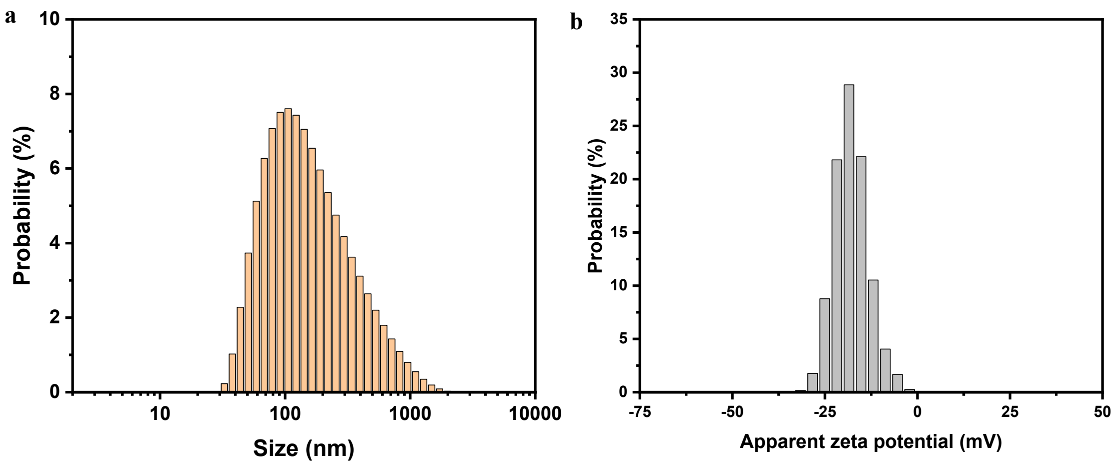


**Figure S2.** (a, b) Distributions of size (a) and zeta potential (b) of liposomal sensors loaded with 100 µM 594-643 in 0.05% NaCl as obtained by dynamic light scattering (DLS) and phase analysis light scattering (PALS), respectively.

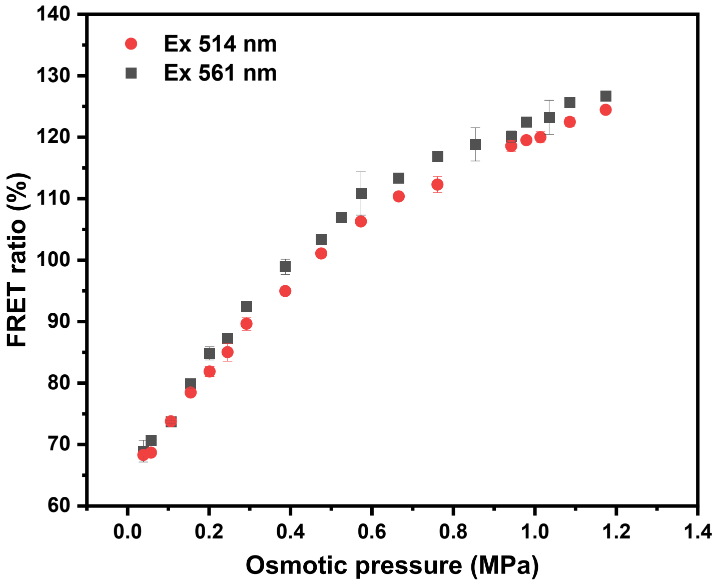


**Figure S3.** FRET ratio *R* obtained with the liposomes loaded with a dye concentration of 50 µM (1:1 molar ratio) in 0.05% NaCl as a function of the external osmotic pressure of NaCl solutions. The excitation wavelength was 514 nm or 561 nm (spectrofluorometer).

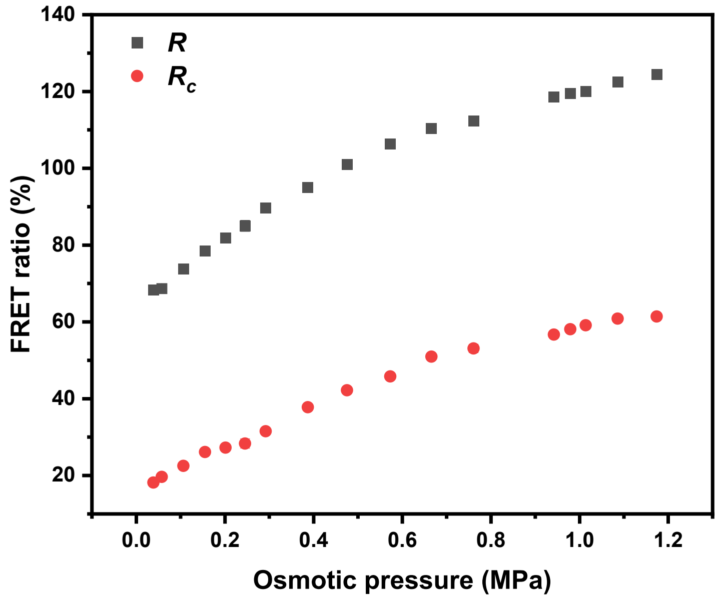


**Figure S4.** FRET ratio *R* obtained with the liposomes loaded with a dye concentration of 50 µM (1:1 molar ratio) in 0.05% NaCl as a function of the external osmotic pressure of NaCl solutions. The excitation wavelength was 514 nm (spectrofluorometer). *R* indicates apparent FRET efficiency and $R_c$ indicates the corrected FRET efficiency.

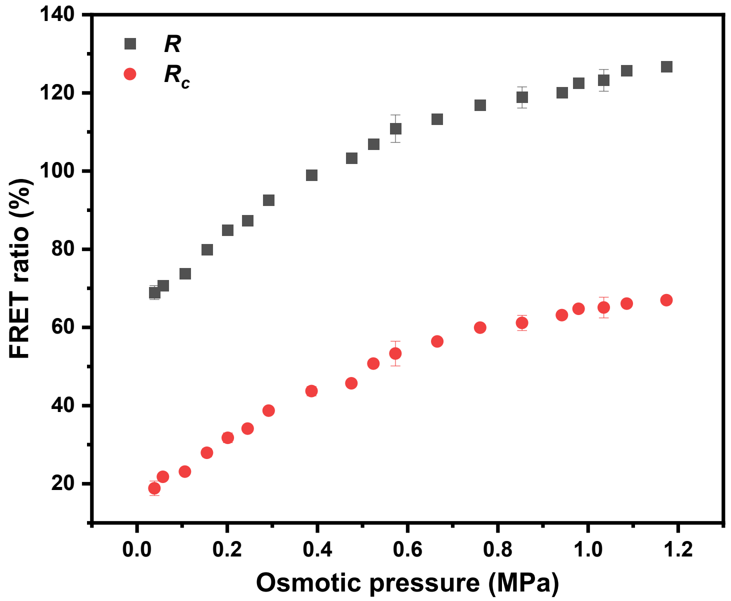


**Figure S5.** FRET ratio *R* obtained with the liposomes loaded with a dye concentration of 50 µM (1:1 molar ratio) in 0.05% NaCl as a function of the external osmotic pressure of NaCl solutions. The excitation wavelength was 561 nm (spectrofluorometer). *R* indicates apparent FRET efficiency and $R_c$ indicates the corrected FRET efficiency.

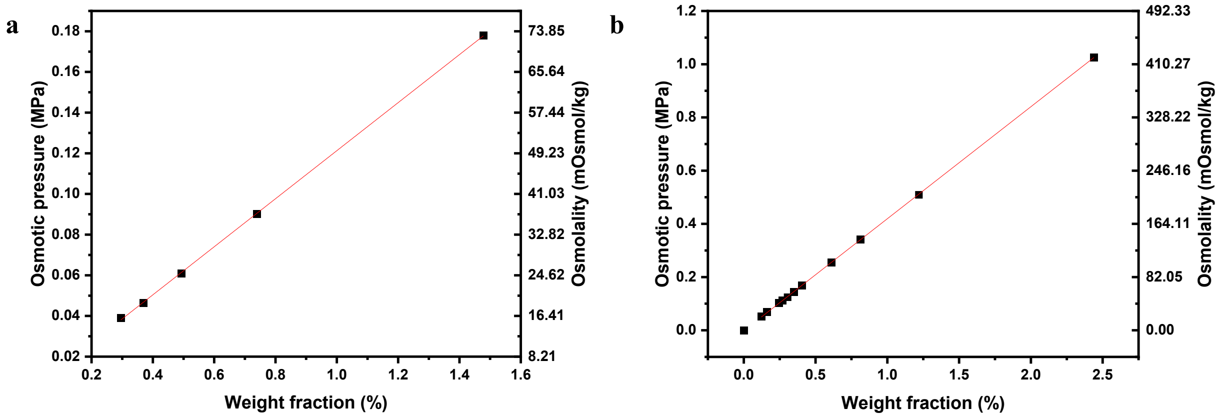


**Figure S6.** Osmolality and osmotic pressure variation of (a) (2% tryptone + 1% yeast extract) (or dilutions) and (b) LB medium (or dilutions) measured with vapor pressure osmometer.

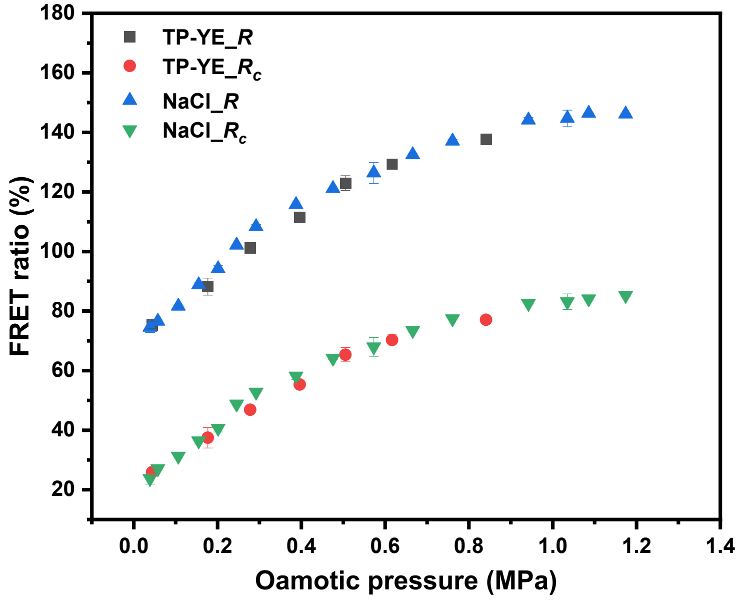


**Figure S7.** FRET ratio $R$ obtained with the liposomes loaded with a dye concentration of 50 µM (1:1 molar ratio) in 0.05% NaCl as a function of the external osmotic pressure of NaCl or tryptone-peptone/yeast-extract/NaCl solutions. The excitation wavelength was 561 nm (spectrofluorometer). $R$ indicates apparent FRET efficiency and $R_c$ indicates the corrected FRET efficiency.

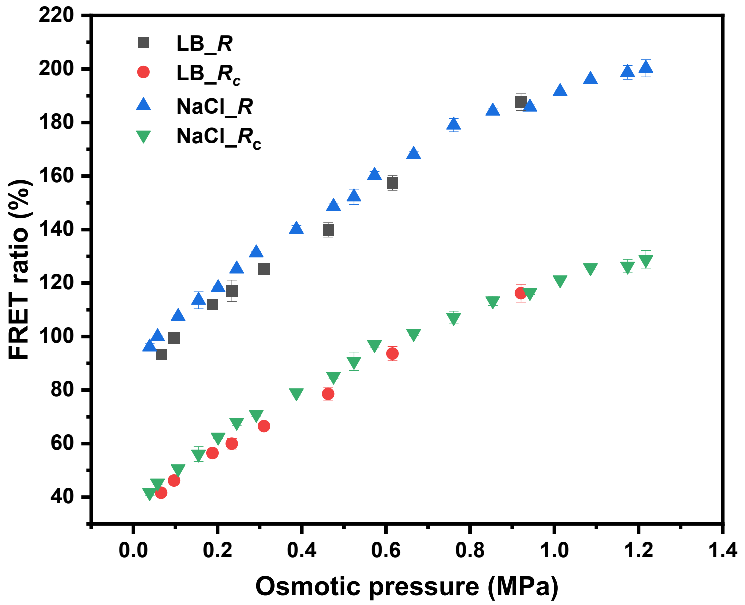


**Figure S8.** FRET ratio $R$ obtained with the liposomes loaded with a dye concentration of 100 μM (1:1 molar ratio) in 0.05% NaCl as a function of the external osmotic pressure of NaCl or LB media and dilutions. The excitation wavelength was 561 nm (spectrofluorometer). $R$ indicates apparent FRET efficiency and $R_c$ indicates the corrected FRET efficiency.

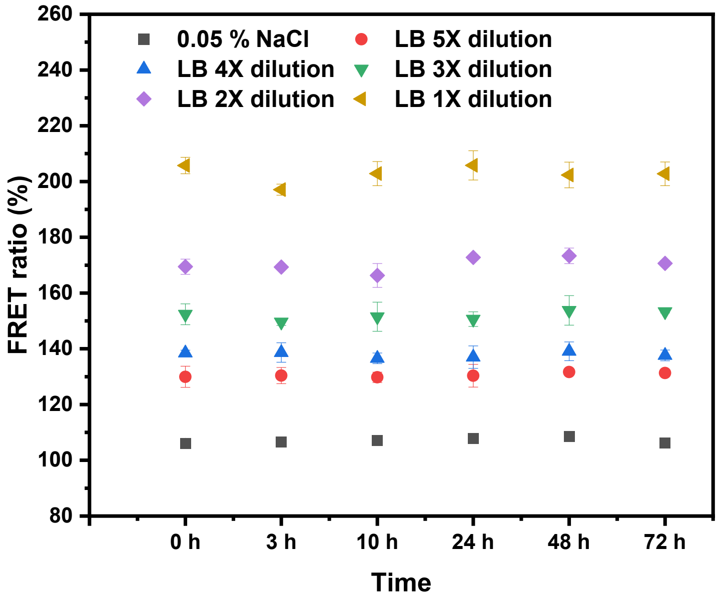


**Figure S9.** FRET ratio $R$ obtained with the liposomes loaded with a dye concentration of 100 μM (1:1 molar ratio) in 0.05% NaCl after incubation in NaCl or LB bacteria culture media and dilutions for 0-72 h at 28 $^{o}$C. The excitation wavelength was 561 nm (spectrofluorometer).

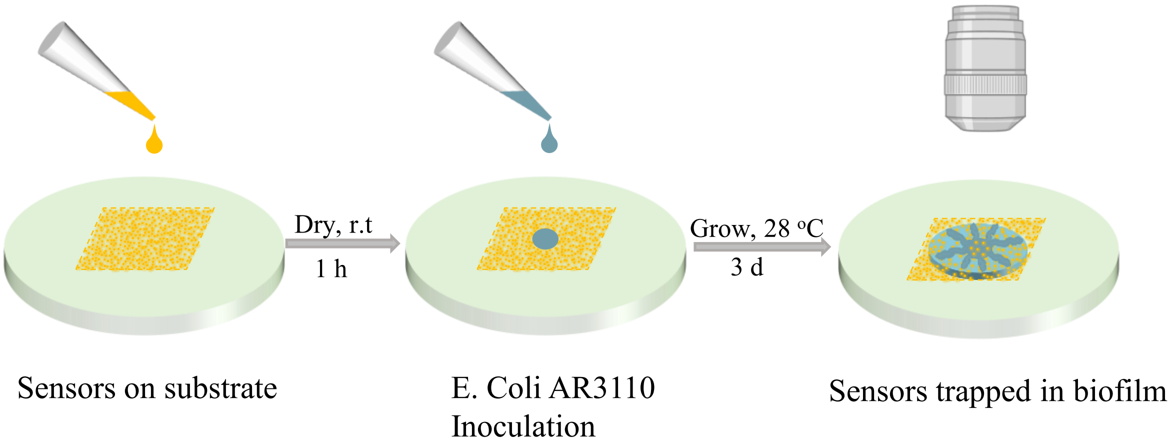


**Figure S10.** Schematic illustration showing the trapping of liposomal sensors in the biofilm growing on agar substrate with nutrients, and the in situ osmotic pressure sensing by using fluorescence microscopy.

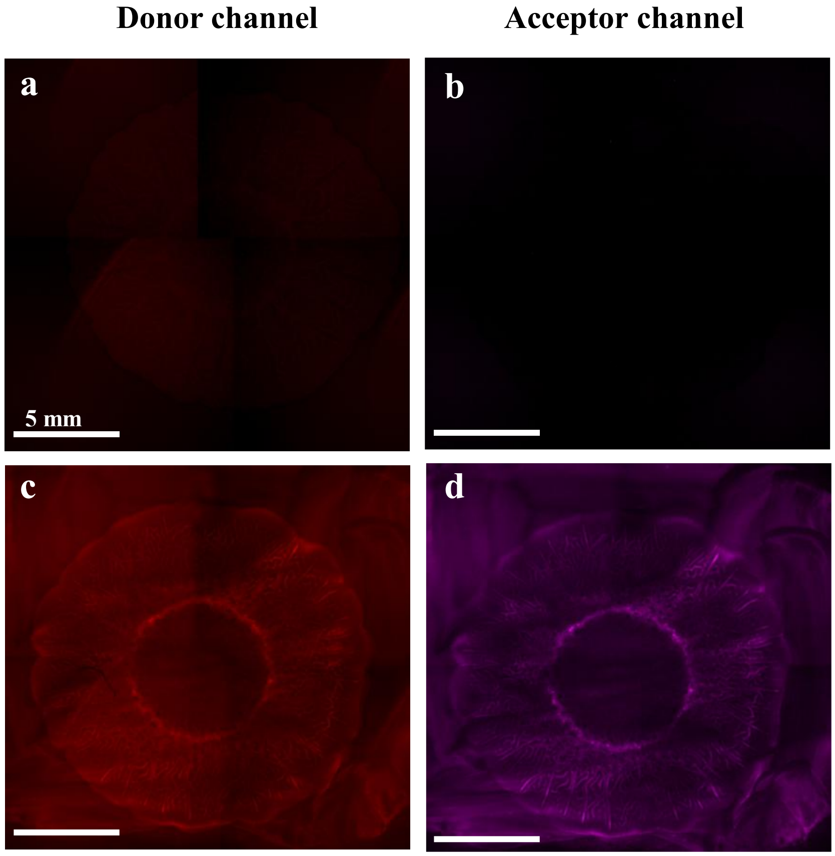


**Figure S11.** (a-d) Wide-field microscopy images of a control *E. coli* biofilm (a, b) and a biofilm with sensors trapped inside (c, d) on the substrate. The fluorescence in (c, d) shows the fluorescence of the sensors in the donor (a, c) and the acceptor (b, d) channel, respectively.

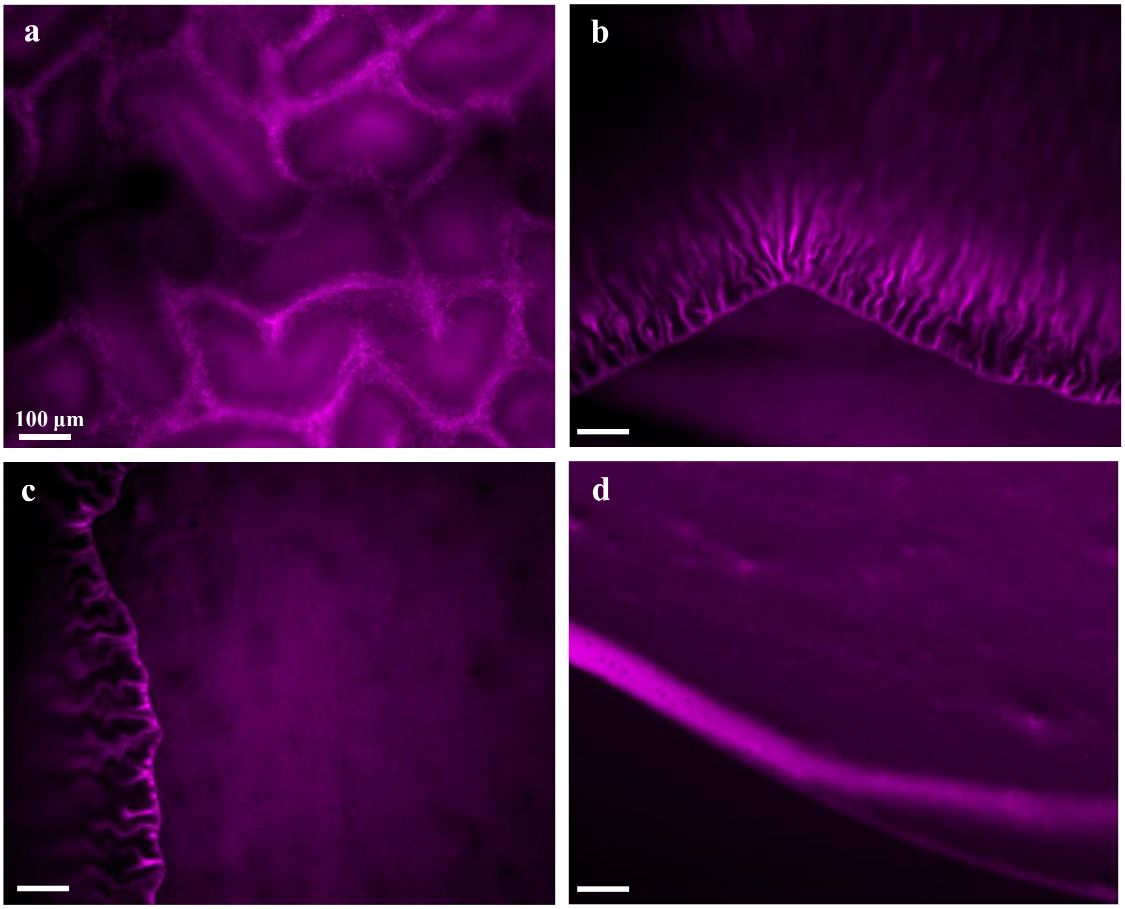


**Figure S12.** (a-d) Wide-field microscopy images of an *E. coli* biofilm with sensors trapped inside on the substrate. The fluorescence shows the fluorescence of the sensors in the acceptor channel in the center (a), periphery (b), and edge (c) region of the biofilm, and in the surrounding substrate (d) region, respectively.

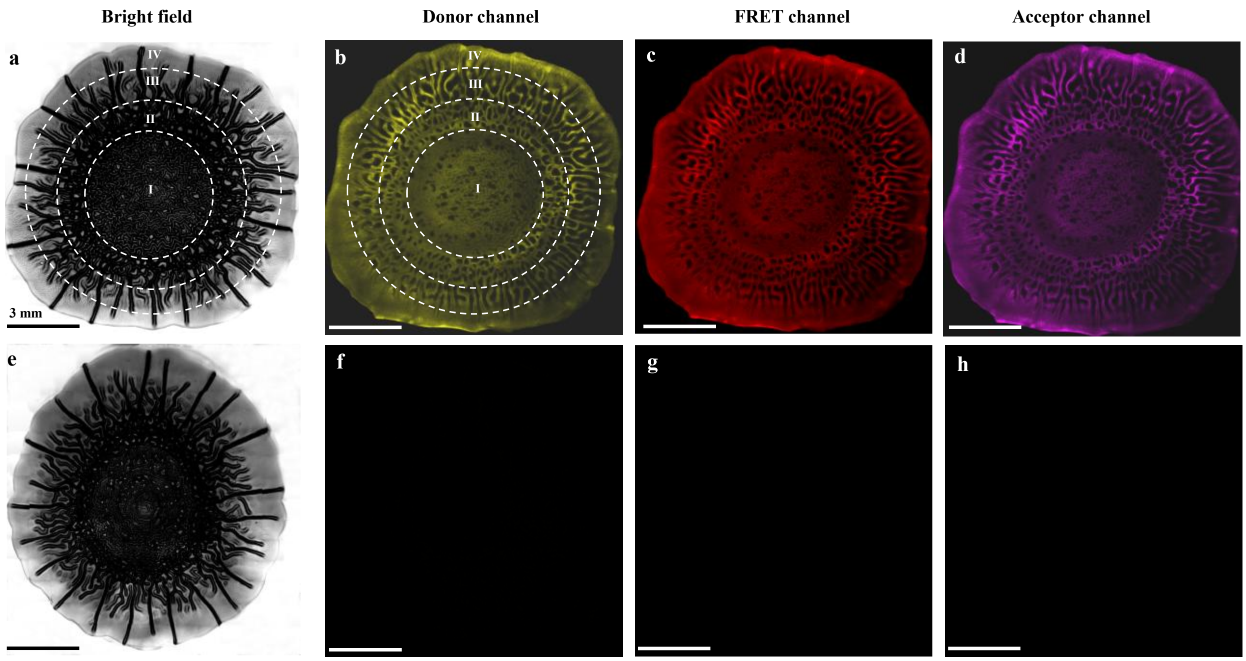


**Figure S13.** Preview CLSM images of a biofilm with sensors trapped inside (a-d) and a control biofilm (e-h). (a,e) Bright field image of the biofilm. (b,f) Donor emission signal (Ex 594 nm, Em 604–620 nm). (c,g) Sensitized acceptor emission signal (Ex 594 nm, Em 680–795 nm). (d,h) Acceptor emission signal (Ex 633 nm, Em 680–795 nm).

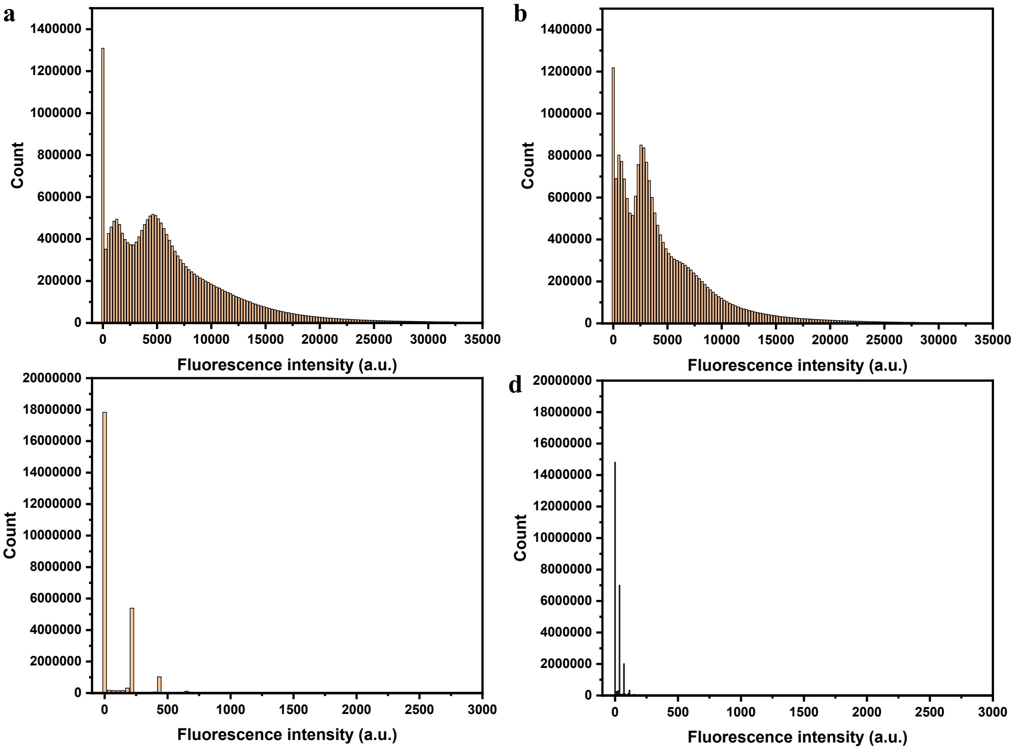


**Figure S14.** Fluorescence intensity histogram of the z-stack of a biofilm with sensors trapped inside (a,b) and a control biofilm (c,d) in the donor channel (a,c; Ex 594 nm, Em 604–620 nm) and the FRET channel (b,d; Ex 594 nm, Em 680–795 nm) measured with CLSM.

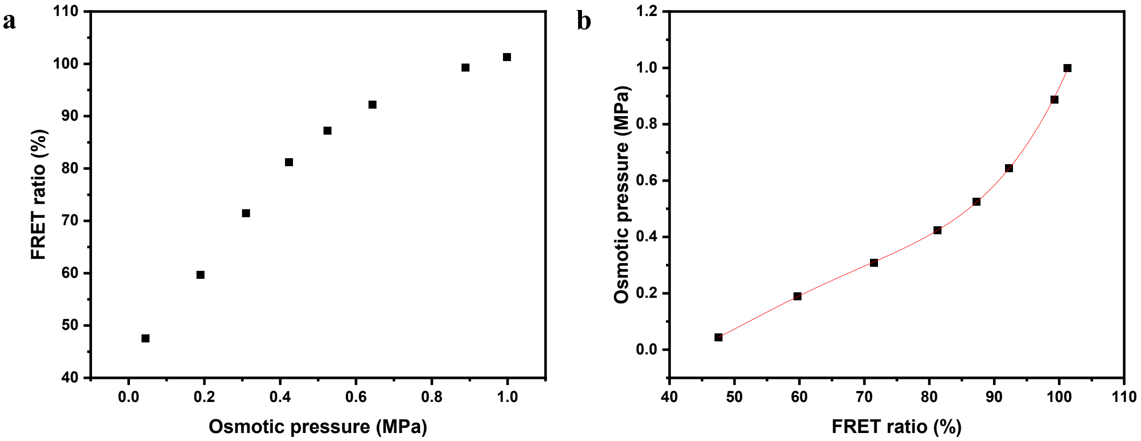


**Figure S15.** Application of sensors for in situ measurement of osmotic pressure in biofilm-substrate system. (a) FRET ratio R obtained with the liposomes loaded with a dye concentration of 100 µM (1:1 molar ratio) in 0.05% NaCl as a function of the external osmotic pressure of tryptone-peptone/yeast-extract/NaCl solutions using CLSM (donor emission signal: Ex 594 nm, Em 604–620 nm; sensitized acceptor emission signal: Ex 594 nm, Em 680–795 nm). (b) Calibration curve used for obtaining Π from R reading. The solid line is an empirical fourth-order polynomial fit to the data points (coefficient of determination = 0.9999).

**Videos S1-S4.** In situ osmotic pressure sensing in the biofilm and in the substrate. (S1-S4) FRET (S1, S3) and osmotic pressure (S2, S4) imaging of the biofilm (z range 150 μm, step 10 μm) (S1, S2) and the substrate (z range 150 μm, step 20 μm) (S3, S4) z-stack from the bottom to the top.
