## Supporting information for "Osmotic pressure gradients in *E. coli* biofilms revealed by in-situ sensors"

### Slide 1
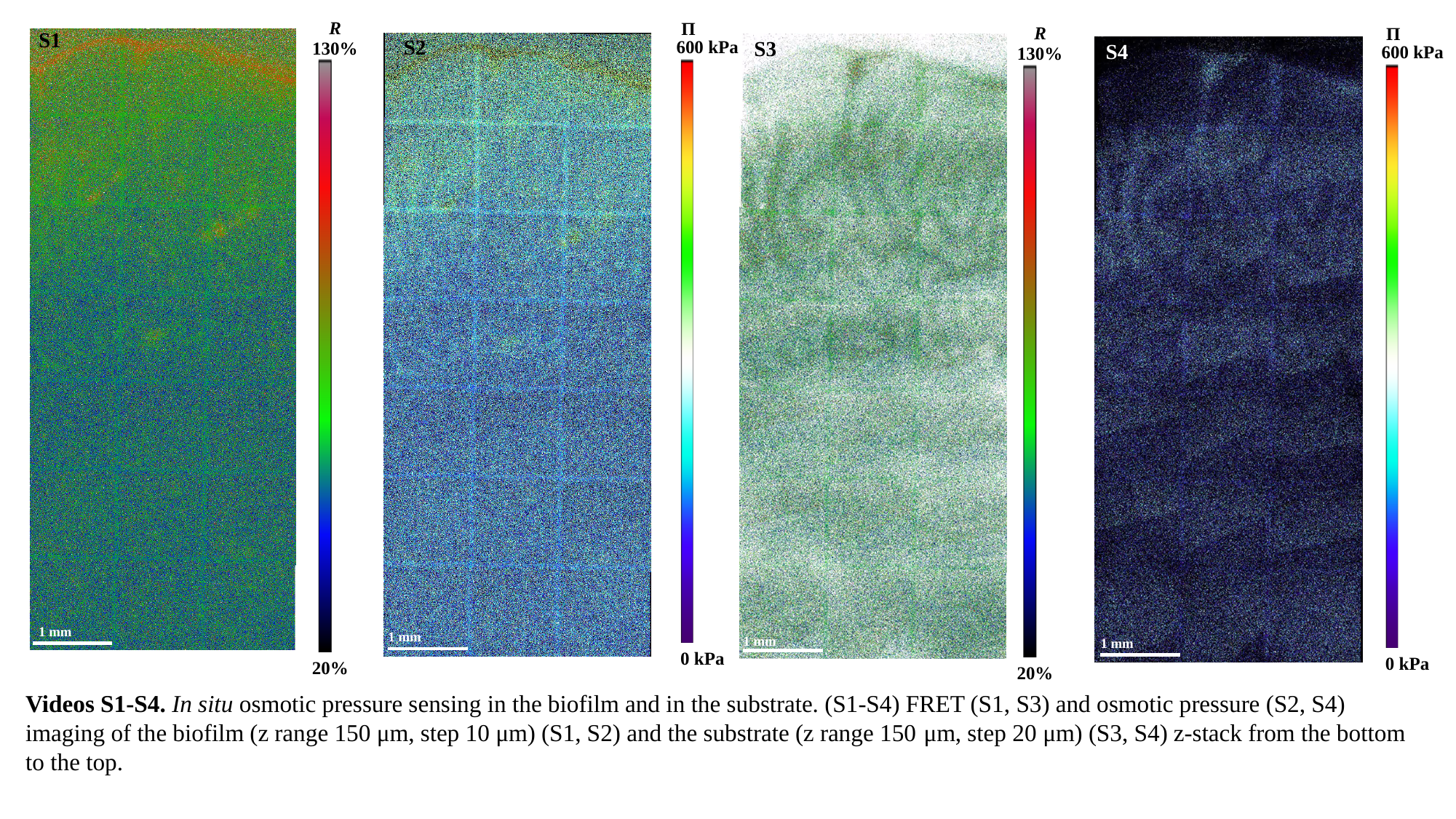

R
130%
20%
П
600 kPa
0 kPa
R
130%
20%
П
600 kPa
0 kPa
S1
1 mm
S2
S3
1 mm
1 mm
S4
1 mm
Videos S1-S4. In situ osmotic pressure sensing in the biofilm and in the substrate. (S1-S4) FRET (S1, S3) and osmotic pressure (S2, S4) imaging of the biofilm (z range 150 μm, step 10 μm) (S1, S2) and the substrate (z range 150 μm, step 20 μm) (S3, S4) z-stack from the bottom to the top.
